# Interferon gamma signaling drives cardiac metabolic rewiring

**DOI:** 10.1101/2025.08.31.673391

**Authors:** Ebram Tharwat Melika, DiyaaElDin Ashour, Kyoungmin Kim, Alexander Nickel, Paula Arias-Loza, Xinyu Chen, Panagiota Arampatzi, Mugdha Srivastava, Martin Väth, Edécio Cunha, Michaela Kuhn, Georg Gasteiger, Ulrich Hofmann, Takahiro Higuchi, Christoph Maack, Stefan Frantz, Anja Karlstaedt, Gustavo Campos Ramos

**Author notes:** Corresponding author: Prof. Dr. Gustavo Ramos, Immunocardiology Laboratory, Comprehensive Heart Failure Centre (CHFC), Am Schwarzenberg 15, 97078 Würzburg, Würzburg, Germany.

## Abstract

**Background:** IFN-gamma (IFN-γ) signaling influences myocardial inflammation and fibrosis across a wide range of conditions, including ischemic and non-ischemic heart failure (HF). However, the direct effects of IFN-γ on cardiomyocytes remain poorly understood. Here, we developed a novel in vivo model to investigate how IFN-γ impacts myocardial metabolism and function.

**Methods:** Male C57BL/6J mice were injected intravenously with hepatotropic adeno-associated virus (AAV2/8) carrying *Ifng* and *nLuc* reporter under the albumin promoter (AAV- *Ifng*) or empty vector control virus (AAV*-*ctrl). Cardiac alterations were monitored on day 28 through flow cytometry, bulk RNA sequencing, targeted metabolomics, isolated mitochondrial activity, echocardiography, and in vivo imaging using [^18^F]fluordeoxyglucose ([^18^F]FDG) and [^18^F]fluoro-6-thia-heptadecanoic acid. Additionally, mice lacking IFN-γ receptor expression in cardiomyocytes (*Myh6*^Cre^ *Ifngr1*^fl/fl^) were used to further dissect the cell-intrinsic roles of IFN-γ signaling in cardiomyocyte metabolic reprograming.

**Results:** After confirming liver-specific viral transfection and elevated serum IFN-γ production at physiological levels, we observed cardiac metabolic adaptation and rewiring in animals treated with AAV-*Ifng* compared to control animals. Myocardial bulk RNA sequencing and gene set enrichment analysis identified an IFN-γ response signature accompanied by marked down-regulations of oxidative phosphorylation and fatty acid oxidation pathways. Functional assessment of isolated cardiac mitochondria showed decreased oxygen consumption, and targeted metabolomics confirmed metabolic shifts toward glycolysis in mice overexpressing IFN-γ. In vivo imaging confirmed increased cardiac glucose uptake following AAV-*Ifng* treatment. Notably, these metabolic alterations were abrogated in mice with cardiomyocyte-specific deletion of IFN-γ receptors (IFNGR).

**Conclusions:** Systemic IFN-γ induces pronounced metabolic reprogramming in the heart, characterized by increased glucose uptake and reduced oxidative phosphorylation, via direct signaling through cardiomyocyte IFNGR. These alterations mirror those observed in aging and some forms of HF, thereby highlighting that, beyond classical inflammation, this cytokine regulates cardiac metabolism.

**Novelty and significance:** *What is known?:*

- Immunological mechanisms can impact myocardial disease progression through complex context-dependent mechanisms.
- IFN-γ, a cytokine primarily secreted by natural killer and T cells, promotes myocardial inflammation and fibrosis in the context of autoimmune myocarditis, pressure-overload-induced heart failure, and Chagas cardiomyopathy.
- Cytokines exert pleiotropic effects and can influence inflammatory responses through mechanisms involving control of energy metabolism.

*What new information does this article contribute?:*

- A novel adeno-associated virus model of systemic IFN-γ elevation allows assessment of cardio-immune-metabolic crosstalk without confounding factors.
- IFN-γ drives cardiac metabolic reprogramming in inflammatory contexts, characterized by enhanced glucose uptake and glycolysis with mitochondrial dysfunction, ultimately altering cardiac metabolic fluxes and function.
- The IFN-γ-induced cardiac metabolic reprogramming is, at least in part, mediated through direct signaling via receptors on cardiomyocytes.

## Introduction

Anti-cytokine-based approaches have regained momentum in cardiology ^1^. Following the success of the CANTOS trial ^2^, which showed reduced cardiovascular risk and heart failure events upon interleukin-1β (IL-1β) neutralization, subsequent studies are now evaluating various anti-cytokine strategies across different myocardial diseases. While most studies have so far focused on targeting pro-inflammatory cytokines, such as IL-1β and IL-6 ^3,4^, that are linked to innate immune mechanisms, emerging evidence suggests that targeting cytokines involved in adaptive immune mechanisms may also be relevant ^5^. Interferon gamma (IFN-γ), a cytokine primarily produced by NK and T cells ^6^, is a key player connecting innate and adaptive immune mechanisms across diverse myocardial pathologies ^7–11^. Further, IFN-γ plays a central role in the development of autoimmune myocarditis ^12,13^ and in Chagas cardiomyopathy, two conditions largely driven by T cell responses ^14^. Moreover, IFN-γ-producing CD4⁺ T cells have been associated with cardiac fibrosis, hypertrophy, and diastolic dysfunction in non-ischemic HF models induced by transverse aortic constriction ^15,16^, whereas IFN-γ signaling is required for effective repair following myocardial infarction (MI) ^17^.

IFN-γ receptors (IFNGR) are broadly expressed across all myocardial cell types, and downstream signaling can influence a wide range of cellular mechanisms. More specifically, IFN-γ signaling promotes macrophage polarization towards a classical inflammatory phenotype through the JAK/STAT1 pathway ^18,19^ and increased expression of major histocompatibility complex in cardiac fibroblasts ^16,20^. However, the direct effects of IFN-γ signaling on cardiomyocytes remain poorly investigated. While its canonical pro-inflammatory impact has been well established across several disease models ^21^, emerging evidence has suggested that IFN-γ can also regulate energy metabolism in targeted cells. For instance, the classical influence of IFN-γ in promoting inflammatory macrophage polarization has been associated with elevated glycolysis and reduced oxidative phosphorylation ^22,23^. In a previous study, we found that T cells undergo spontaneous clonal expansions in the heart and draining lymph nodes during healthy aging, characterized by marked increase in IFN-γ production. Moreover, we found that this age-related increase in myocardial IFN-γ response signature was linked to altered expression of several genes involved in myocardial metabolism, including oxidative phosphorylation, fatty acid metabolism, and glycolysis ^24^. Previous in vitro studies also reported IFN-γ-mediated effects on mitochondrial homeostasis in cultured cardiomyocytes ^25^. These observations raise the possibility that IFN-γ may modulate cardiac metabolic processes and function in vivo, thereby contributing to HF development through mechanisms beyond its well-established inflammatory roles.

To mechanistically address this question, we devised an adeno-associated virus (AAV)- based model to induce chronic systemic IFN-γ expression, thus enabling us to dissect its metabolic impacts on the myocardium in the absence of concomitant myocardial disease and confounding factors. Building on this model, our study integrates immune-phenotyping, bulk RNA sequencing, targeted metabolomics, assessment of mitochondria function, and in vivo positron emission tomography (PET) to evaluate cardiac glucose and fatty acid uptake in response to IFN-γ overexpression. Our findings reveal that systemic IFN-γ elevation induces cardiac metabolic rewiring marked by reduced oxidative phosphorylation, reminiscent of some HF models ^26^. Moreover, by studying cardiomyocyte-specific IFNGR1 knockout mice, we demonstrated that direct IFN-γ signaling is sufficient to induce cardiac metabolic in cardiomyocytes. Taken together, these results shed new light on the roles of a key cytokine in the myocardium and demonstrate novel mechanisms of cardio-immune-metabolic regulation.

## Methods

The full methods and supplemental figures are available in the Supplemental Material.

### Data availability

The transcriptomic data acquired in this study have been deposited in NCBI’s Gene Expression Omnibus (GEO) database (GSE303910).

### Study approval

All animal protocols were approved by the local authorities (Regierung von Unterfranken, Würzburg, Germany) and the experiments were performed according to the Federation for Laboratory Animal Science Associations (FELASA) guidelines ^27^.

### Statistical analyses

The results are shown as the mean ± the standard deviation of the mean (SD) along with the distribution of all individual values in each group. Sample sizes for each group are described in figure legends. Graphs and statistical analyses were performed with GraphPad Prism (version 7.0, GraphPad Software, San Diego, CA, USA). Unpaired two-tailed t-test was used to compare two groups with data following normal distribution. For multiple comparisons between more than two groups, one- or two-way analyses of variance (ANOVA) were conducted followed by post hoc test. Differences were considered significant for p values below 0.05.

## Results

### Establishing an experimental model to investigate IFN-γ impacts on myocardial biology

To mechanistically dissect how IFN-γ impacts cardiac metabolism in the absence of concomitant myocardial disease and confounding factors, we generated a hepatotropic adeno-associated virus (AAV), serotype 2/8 to induce liver-specific IFN-γ expression, along with a NanoLuc (NLuc) reporter under the albumin promoter (AAV-*Ifng)*. As a control, we used AAV2/8 carrying only the reporter gene (AAV-ctrl). Initial in vitro validation in AML12 hepatocytes confirmed a dose-dependent increase in luciferase activity and IFN-γ secretion in response to AAV-*Ifng,* but not AAV-ctrl, treatment (Supp. Fig. 1). Next, C57BL/6J male mice were intravenously injected with 10¹¹ viral genomes (vg) per mouse of AAV-*Ifng* or AAV-ctrl to produce systemic IFN-γ overexpression in healthy mice, and endpoint analyses were performed 28 days post-injection (**Fig. 1A**). Specific viral transduction was confirmed by evaluating luciferase activity across various organs and detecting it exclusively in the liver (**Fig. 1B**). *Ifng* mRNA expression was specifically augmented in livers of AAV-*Ifng* mice but not in other tissues or in mice injected with AAV-control. Further, AAV-*Ifng*-treated mice showed elevated circulating IFN-γ levels comparable to those observed in mice infected with acute lymphocytic choriomeningitis virus (**Fig. 1B**), whereas AAV-ctrl-treated mice had no rise in plasma IFN-γ levels. Western blot analysis revealed heightened phospho-STAT1 levels that demonstrated signaling in liver and heart of AAV-*Ifng* mice, but not controls (**Fig. 1C**). These findings confirm that AAV-*Ifng* administration successfully generated liver-specific IFN-γ expression, resulting in higher circulating IFN-γ levels within a physiologically relevant range, sufficient to trigger both local and systemic signal transduction.

**Figure 1:**
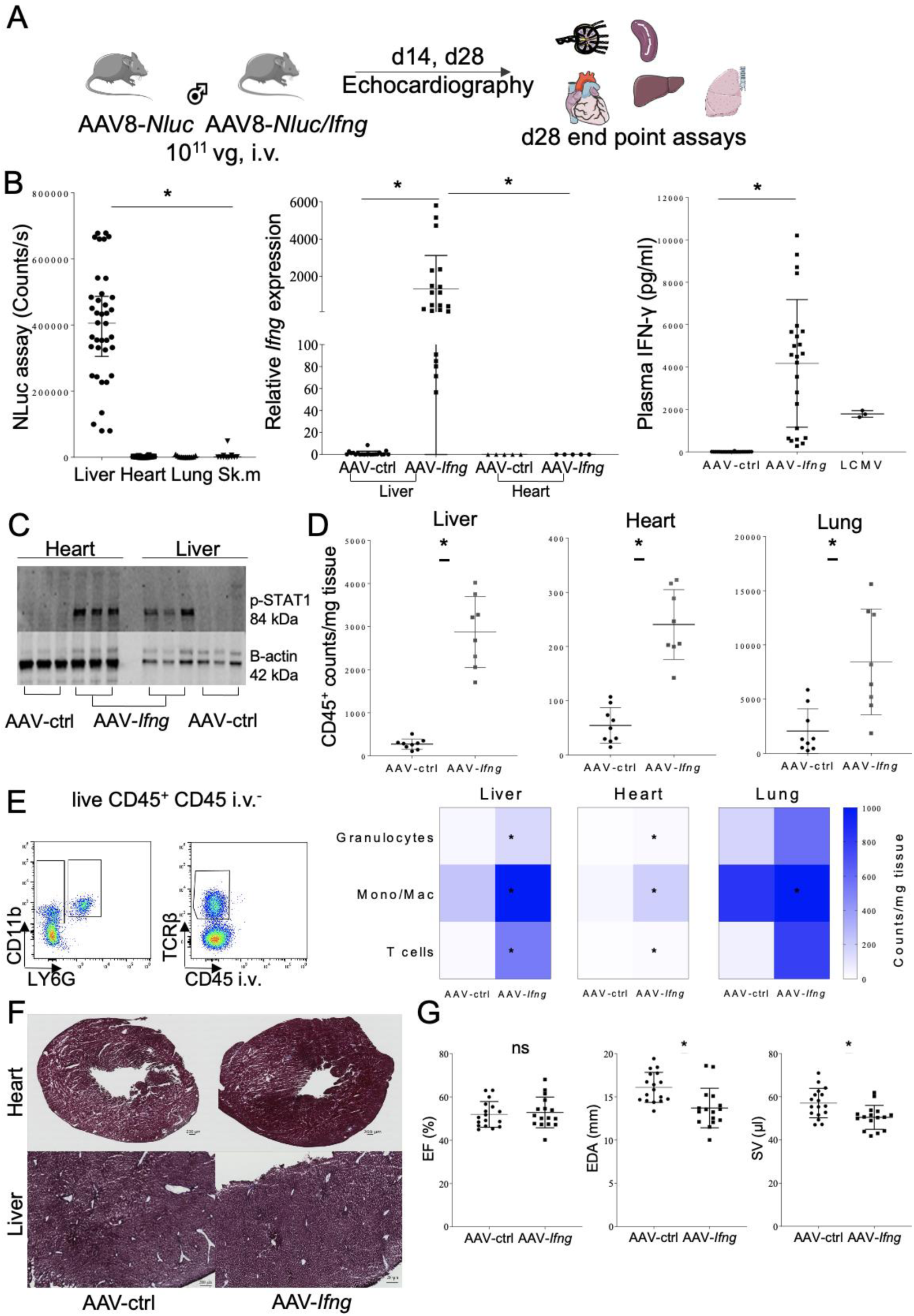
Systemic IFN-γ expression induces myocardial alterations. **A**: Schematic representation of the in vivo mouse model. **B**: Luciferase activity in liver, heart, lung, and skeletal muscles; relative *Ifng* expression in liver and heart of AAV-*Ifng* injected mice versus AAV-*Nluc*-injected control mice; and IFN-γ plasma concentration compared between the two groups and LCMV-injected mice. **C**: Western blot membrane showing p-STAT1 and β-actin in livers and hearts of the two groups (3 different mice per group). **D**: CD45^+^ cells / mg tissue analyzed via flow cytometry, comparing different organs from the two groups. **E**: Flow cytometry gating to distinguish granulocytes (CD45^+^ Cd11b^+^ Ly6G^+^), monocytes and macrophages (CD45^+^ Cd11b^+^ Ly6G^-^), and T cells (CD45^+^ TCRβ^+^) with heat map quantifications of different cell populations in liver, heart, and lung, compared between the two groups. **F**: Representative Masson’s trichrome staining of heart and liver tissues from the two groups (3 different mice per group). **G**: Day 28 echocardiographic findings indicating EF (%), EDA (mm), and SV (µl). Statistical analysis using t-test. * Indicates significance with p value < 0.05. Graphs B, D, G are scatter plots presenting Mean ± SD.

Flow cytometry analysis of enzymatically dissociated tissues indicated that AAV-*Ifng* administration induced mild systemic inflammation with increased CD45^+^ immune cell counts / mg tissue in the liver, heart, and lungs (**Fig. 1 D**). Additionally, IFN-γ over-expression mildly increased counts of tissue granulocytes (CD45^+^ CD11b^+^ Ly6G^+^), T cells (CD45^+^ CD11b^-^ TCRβ^+^), and especially monocyte/ macrophages (CD45^+^ CD11b^+^ Ly6G^-^) (**Fig. 1 E, Supp. Fig. 2**). Moreover, significant expansions of dendritic cells (CD11c^+^ MHCII^+^), macrophages (CD64^+^), monocyte-derived macrophages (CD64^+^ CCR2^+^), and pro-inflammatory monocytes (CD64^-^ Ly6C^+^) were observed in heart, liver, and lungs of AAV-*Ifng* injected mice (**Supp. Fig. 2 A, B**). AAV-*Ifng* administration also impacted T cell distributions and phenotypes across various organs (**Supp. Fig. 2 C, D, E**). Additionally, we noted IFN-γ-dependent up-regulation of programmed death ligand 1 (PD-L1) and major histocompatibility II (MHCII) on endothelial cells (CD45^-^ CD31^+^) from the liver, heart, and lung (**Supp. Fig. 2 F, G**). Despite the low-grade systemic inflammation produced by AAV-*Ifng* treatment, no gross morphological alterations were found in cardiac and hepatic histological slices stained with Masson’s trichrome (**Fig.1 F**). In contrast to other models of constitutive IFN-γ expression ^28^, the absence of major tissue injury and weight loss in our model supports its physiological relevance, in line with previous observations during healthy aging and in models of HF ^24,29,30^. Finally, echocardiographic analysis performed on day 28 post-infection revealed a significant IFN-γ- dependent reduction in end-diastolic area and stroke volume, with preserved ejection fraction in AAV-*Ifng-*injected mice suggesting that chronically elevated IFN-γ levels can drive cardiac functional impairment (**Fig. 1 G**).

### Systemic IFN-γ induces transcriptional alterations in pathways associated with myocardial metabolism

To further characterize the cardiac alterations induced by systemic IFN-γ, we performed bulk RNA sequencing on myocardial and liver samples. Gene set enrichment analysis (GSEA) in myocardial samples confirmed that the ‘IFN-γ response’, and ‘inflammatory response’ gene sets were among the top up-regulated in AAV-*Ifng-*treated mice, thus supporting the validity of this model. Interestingly, ‘oxidative phosphorylation’, ‘fatty acid oxidation’, and ‘mitochondrial respiratory chain’ were among the top down-regulated gene sets in the heart and liver of AAV-*Ifng-*treated mice (**Fig. 2, Supp. Fig. 3, Supp. Table 1**). These changes were partially driven by decreased expression of *Idh3a*, *Timm9*, *Pdha1*, and *Atp5b,* which are encoding for key-regulatory enzymes of the respiratory chain complexes and Krebs cycle. Furthermore, AAV-*Ifng* treatment significantly up-regulated genes linked to glycolysis, including *Ppfia4*, *G6pd*, *Pfkp*, and *Pdk3* (**Fig. 2 C)**. Conversely, the ‘fatty acid oxidation’ gene set signature showed down-regulation of *Mgll*, *Acsl1*, and *Crat* transcripts (**Fig. 2 C**). Overall, these observations suggest a metabolic shift from fatty acid oxidation towards glycolysis in response to IFN-γ signaling in the heart.

**Figure 2:**
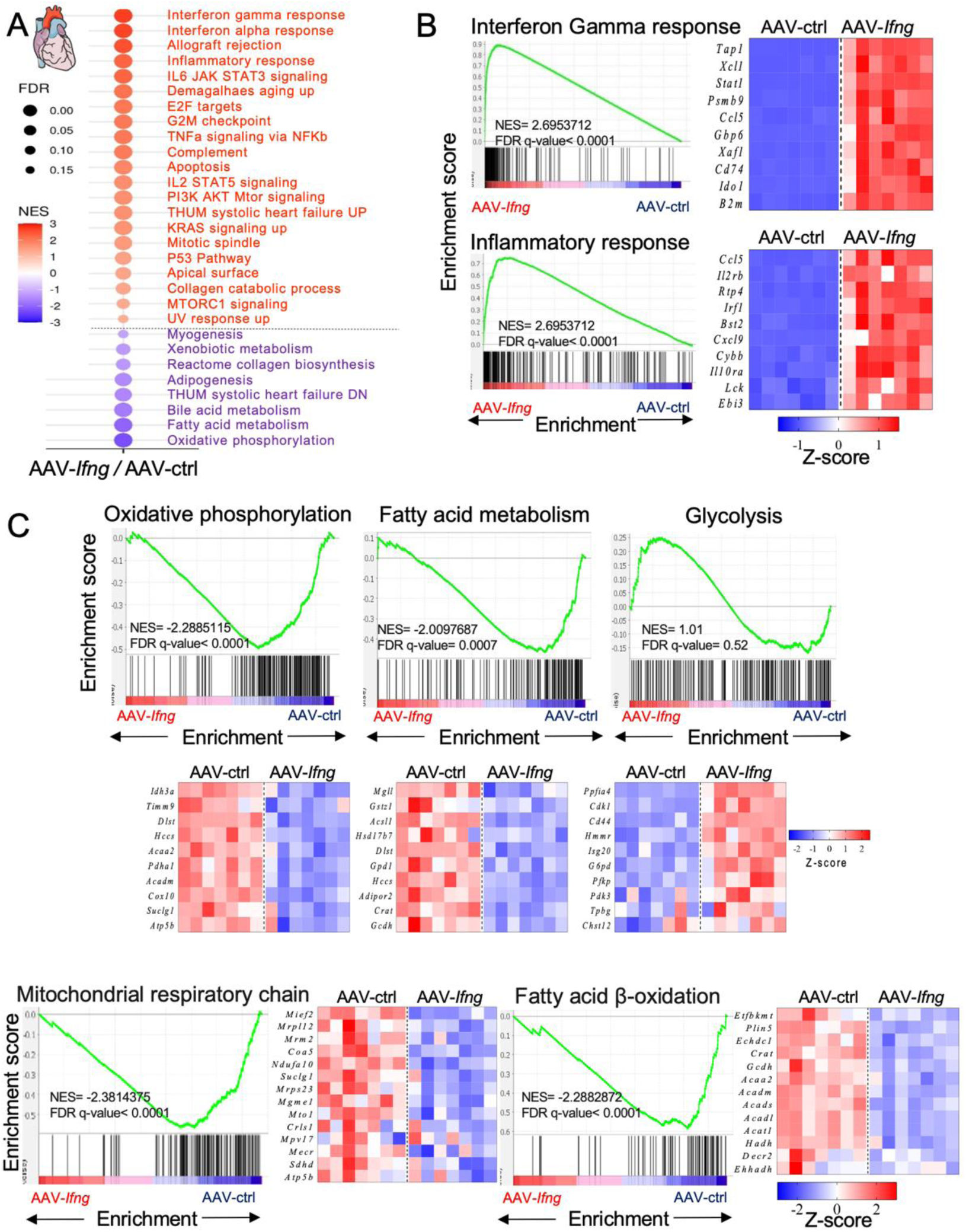
IFN-γ causes transcriptional alterations in pathways linked to cardiac metabolism. **A**: Dot blot showing significant gene set up- and down-regulation as obtained from bulk RNA sequencing data of hearts from the two groups. Color scale represents the normalized enrichment score, and circle size shows the false discovery rate (FDR). **B**: Gene set enrichment analysis of interferon gamma response and inflammatory response, with heat maps of z-scores for representative top genes, in the heart, among the two groups. **C**: Gene set enrichment analysis of oxidative phosphorylation, fatty acid oxidation, glycolysis, mitochondrial respiratory chain, and fatty acid β-oxidation with heat map of z-scores for representative top genes, in the heart, among the two groups. Blue denotes transcript down-regulation while red indicates up-regulation.

### IFN-γ promotes cardiac metabolic rewiring

Transcriptional alterations prompted us to investigate in further detail how IFN-γ signaling could induce cardiac metabolic rewiring. To assess whether transcriptional shifts translated into changes in protein expression and overall metabolism, we conducted gas chromatography and mass spectrometry-based metabolomics on freeze-clamped hearts, measured key enzymatic activities, and functionally characterized isolated mitochondria (**Fig. 3**). Initial principal component analysis of 61 cardiac metabolites revealed clear differences between AAV-*Ifng*- and AAV-ctrl-injected mice (**Fig. 3 A**). Among the metabolites investigated, glucose; lactate; and amino acids such as alanine, glutamate, aspartate, and succinate were significantly increased in the hearts of AAV-*Ifng* injected mice, suggesting altered glucose utilization, enhanced amino acid metabolism, and disrupted Krebs cycle. Moreover, cholesterol and glycerol were also significantly elevated, potentially indicating lipid metabolism remodeling (**Fig. 3 B**). We found sustained lactate dehydrogenase (LDH) activity in heart tissue from AAV-*Ifng* mice compared to controls, while pyruvate dehydrogenase activity was significantly reduced (**Fig. 3 C**). These findings are corroborated by RNA-seq showing down-regulation of *Pdha1* and up-regulation of *Pdk3*, which phosphorylates and inhibits PDH. Western blotting confirmed reduced protein expression of ATP synthase beta (ATP5B) (**Supp. Fig. 4 A**), a subunit of complex V within the electron transport chain. In contrast, glyceraldehyde 3-phosphate dehydrogenase (GAPDH) expression, a key metabolic enzyme within glycolysis, increased in heart tissue samples from AAV-*Ifng* mice. No differences were observed in the expression of the mitochondrial voltage-dependent anionic channel (VDAC) (**Supp. Fig. 4 A**), which is a key mitochondrial outer membrane protein and regulator of membrane permeability ^31^.

**Figure 3:**
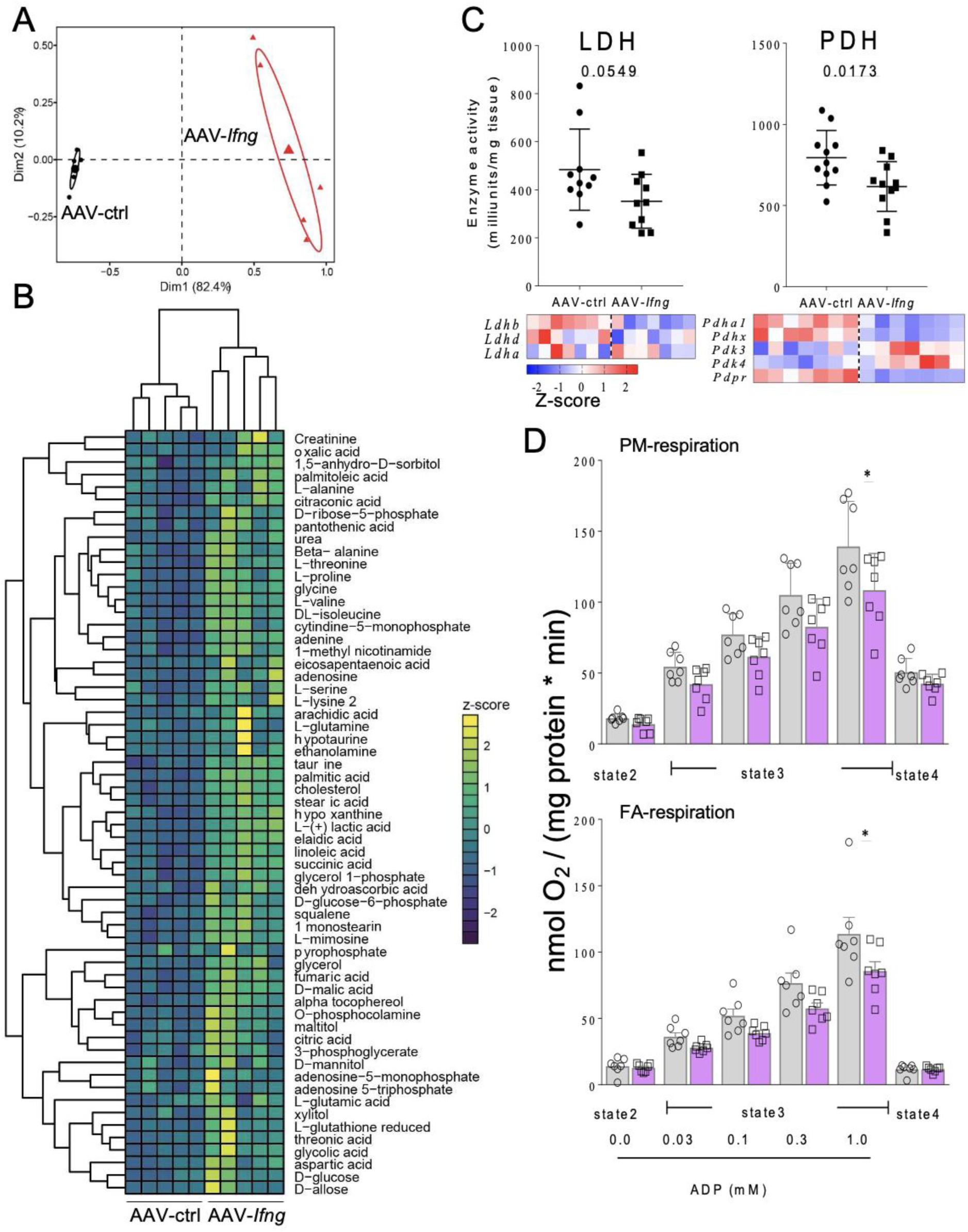
IFN-γ signaling causes cardiac metabolic reprogramming. **A**: Principal component analysis of the cardiac metabolome of AAV-*Ifng*- and AAV-ctrl-injected mice (n=5 per group). **B**: Heatmap representing unsupervised hierarchical cluster analysis of the cardiac metabolite profile of each mouse. **C**: Lactate dehydrogenase enzyme activity quantification graph comparing the two groups, heatmap of z-scores for enzyme genes, pyruvate dehydrogenase enzyme activity quantification graph comparing the two groups, and heatmap of z-scores for enzyme genes. **D**: Quantification of oxygen consumption in both pyruvate malate (PM)- and palmitate (fatty acid, FA)- complex-dependent respiration in the absence (state 2) or presence of increasing ADP concentrations (state 3), and in response to the ATP synthase inhibitor oligomycin (state 4). Statistical analysis used t-test. * Indicates significance with p value < 0.05. Graphs C, D are scatterplots showing Mean ± SD. Heat maps of z-scores for representative top genes in the hearts of the two groups. Blue represents transcript shows down-regulation while red indicates up-regulation.

Functional analyses of isolated cardiac mitochondria further confirmed that IFN-γ expression blunted oxygen consumption rates during both pyruvate/malate (PM)- and palmitate-dependent, but not succinate respiration under state 3 conditions (i.e., in the presence of saturating concentrations of ADP; **Fig. 3D** and **Supp. Fig. 4B**). Mitochondrial H_2_O_2_ emission or membrane potential (ΔΨ_m_) were not different during PM- or fatty acid (FA)-coupled respiration in the absence or presence of ADP, while H_2_O_2_ emission was slightly decreased in the AAV-*Ifng* group in the presence of antimycin A, an inhibitor of complex III which provokes maximal superoxide formation as a positive control (**Supp. Fig. 4C** and D). Taken together, these findings verify that IFN-γ overexpression rewires cardiac metabolism at multiple levels, including transcription, protein expression, enzyme activity, and mitochondrial function, ultimately leading to impaired oxidative metabolism and a shift toward glycolytic and amino acid-driven energy pathways.

### Metabolic flux modeling reveals pathways preferentially impacted by IFN-γ signaling

To further interrogate these observations at a systems level, we integrated our multi-omics data (RNA-seq, metabolomics, enzyme activities) into metabolic flux analysis using CardioNet ^32–36^, a genome-scale mammalian heart-specific metabolic network. We calculated flux rates for AAV-ctrl and AAV-*Ifng* mice with experimental measurements as constraints for enzymes and metabolites represented in CardioNet. Using these constraints, we determined flux rates for each network reaction using computational modelling by flux balance analysis (FBA), which applies a steady-state assumption while optimizing an objective function ^32,34,36,37^. Optimal flux solutions are estimated based on the linear optimization problem while fulfilling experimental constraints defined by experimental constraints. Unsupervised hierarchical cluster analysis of significantly altered flux rates (**Fig. 4A** and **Supp. Fig. 5**) demonstrated a distinct metabolic remodeling between AAV-ctrl and AAV-*Ifng* mice. Consistent with our RNA-seq and metabolomics analysis, CardioNet simulations revealed that increased IFN-γ signaling in the heart promotes glucose uptake and glycolytic flux (**Fig. 4B**). Increased glucose uptake is predicted to enhance flux through the oxidative pentose phosphate pathway and increase glucose oxidation via the Krebs cycle. In addition, the modelling revealed an increased contribution of glutamine-based anaplerosis of Krebs cycle intermediates to maintain oxidative phosphorylation and energy provision. Intriguingly, the modelling predicted an increased cleavage of citrate into oxaloacetate and acetyl-CoA via the ATP-dependent citrate lyase resulting in cholesterol biosynthesis. Taken together, FBA reveals an IFN-γ dependent metabolic flux adaptation, which increases flux through glycolysis and pentose phosphate pathway and shifts cardiac energy provision towards glucose and amino acids.

**Figure 4:**
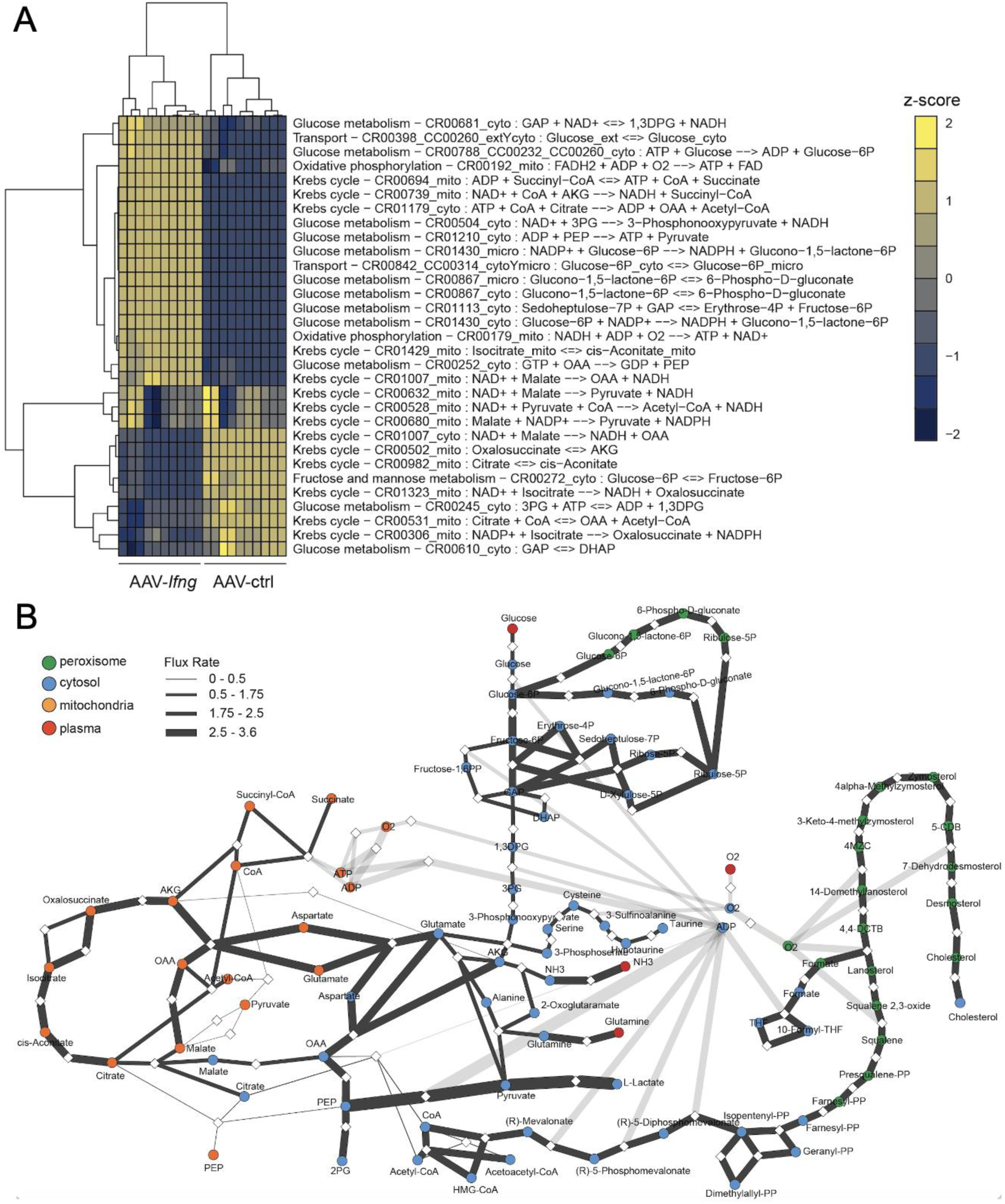
Metabolic flux modeling reveals pathways preferentially impacted by IFN-γ signaling. **A**: Heatmap representing unsupervised hierarchical cluster analysis of flux balance analysis with an adjusted p-value<0.01. Experimental data from AAV-ctrl- and AAV-*Ifng*-injected mice was integrated into CardioNet simulations, and metabolic flux rates were calculated using an objective function demanding energy provision and biomass synthesis. Calculated flux rates were compared using a linear regression model and annotated to CardioNet pathways. **B**: Comparative metabolic flux analysis of AAV-ctrl- and AAV-*Ifng*-injected mice using CardioNet-based simulations. Network graph is constructed from flux balance solutions. Metabolites and enzymes are depicted as circles and diamonds, respectively. Reactions are assigned to cytosolic, mitochondrial, peroxisomal or extracellular metabolic fluxes. The thickness of each line connecting metabolites and proteins indicates the calculated flux rate in AAV-*Ifng*-injected mice compared to AAV-ctrl. Abbreviations: 4MZC, 4alpha-Methylzymosterol-4-carboxylate; 4,4-DCTB, 4,4-Dimethyl-5alpha-cholesta- 8,14,24-trien-3beta-ol; 5-CDB, 5alpha-Cholesta-7,24-dien-3beta-ol.

### In vivo PET imaging reveals increased cardiac glucose uptake in response to systemic IFN-γ stimulation

Our experimental and computational analysis revealed a metabolic substrate switch from fatty acids to glucose upon overactivation of IFN-γ in the heart. To validate the computational predictions, we quantified cardiac [^18^F]-fluordeoxyglucose ([^18^F]-FDG) and [^18^F]fluoro-6-thia-heptadecanoic acid ([^18^F]-FTHA) fatty acid uptake in vivo following AAV-ctrl and AAV-*Ifng* treatment. [^18^F]-FDG uptake was significantly higher in the hearts of the AAV-*Ifng* group compared to AAV-*ctrl* (**Fig. 5 A, B; Supp. Fig. 6 A**), confirming an increased glucose uptake upon IFN-γ signaling. We did not observe changes in systemic glucose and insulin levels at fasting (**Supp Fig. 6 B**). Intriguingly, we observed a positive linear correlation between glucose uptake and IFN-γ serum levels (**Fig. 5 C**). These findings align with our computational modelling and multi-omics data (metabolomics and RNA-seq), which show IFN-γ-dependent up-regulation of *Slc2a1*, *Slc2a6*, and *Slc2a9* (which encode for glucose transporter 1 (GLUT1), GLUT6, and GLUT9, respectively) (**Fig. 5 D**). Surprisingly, [^18^F]-FTHA uptake remained unaltered in the heart, but was significantly reduced in the liver upon AAV- *Ifng* treatment (**Fig. 5 E**). Plasma triglycerides levels were not changed between groups (**Supp. Fig. 6 C**), indicating that AAV-*Ifng* impairs fatty acid uptake in the liver. Taken together, in vivo PET imaging corroborates an IFN-γ-dependent switch towards increased glucose uptake in the heart.

**Figure 5:**
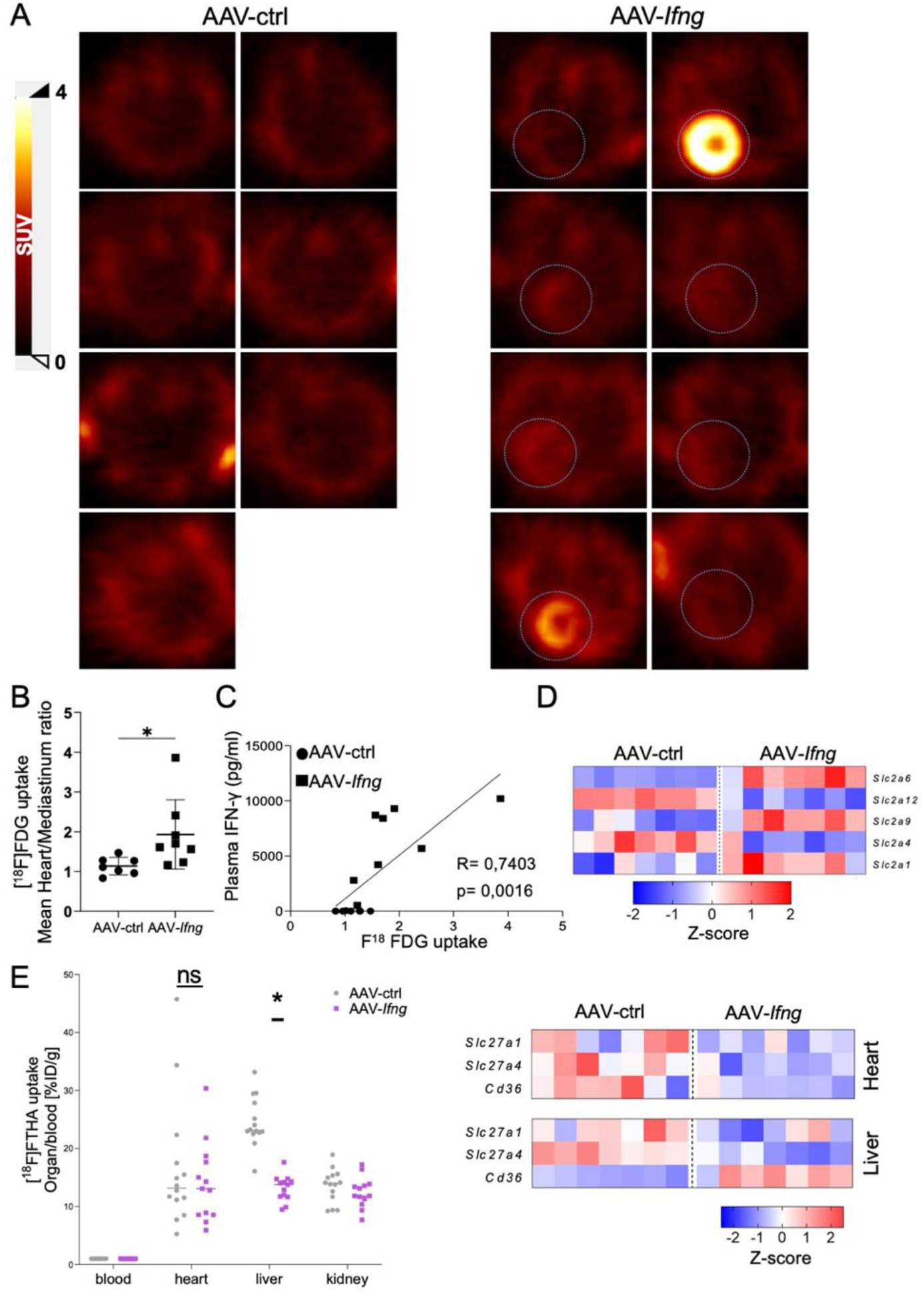
PET imaging reveals glucose and fatty acid uptake in the heart. **A**: Representative [^18^F]FDG glucose uptake PET images from the transverse plane in both AAV- ctrl- and AAV-*Ifng*-injected mice. **B**: Quantification of mean [^18^F]FDG glucose uptake heart/mediastinum ratio in AAV-*Ifng*-injected mice compared to controls. **C**: Linear regression correlation between IFN-γ serum level and [^18^F]FDG glucose uptake. **D**: Heatmap of z-scores for glucose transporter genes, in the heart, among the two groups. **E**: Quantification of [^18^F]FTHA biodistribution across different organs in AAV-*Ifng*-injected mice compared to controls, with (on right) heat maps of z-scores for fatty acid transporters in both heart and liver, comparing the two groups. * Indicates significance with p value < 0.05. Graphs A, B, C, D are scatterplots showing mean ± SD. Heat maps of z-scores for representative top genes, in the heart, among the two groups. Blue colour represents transcript down-regulation while red indicates up-regulation.

### IFN-γ-induced cardiac metabolic rewiring is driven by direct effects in cardiomyocytes

Since IFNGRs are broadly expressed across different cell populations in the myocardium, we sought to identify the cell types primarily linked to the observed metabolic effects. Myocardial immunofluorescence staining of mice treated with AAV-*Ifng* revealed widespread p-STAT1 distribution in both cardiomyocytes (Phalloidin^+^ cells) and non-cardiomyocytes (Phalloidin^-^ cells) (**Fig. 6 A**). Thus, to dissect whether the direct impact of IFN-γ on cardiomyocytes drives the demonstrated cardiac metabolic rewiring, we crossed *Myh6*^Cre^ and *Ifngr1*^fl/fl^ mice to generate animals lacking IFNGR specifically in cardiomyocytes (IFNGR^CM-KO^) (**Fig. 6 B**). IFNGR^CM-KO^ exhibited significantly reduced myocardial but not liver *Ifngr1* expression, compared to controls (**Supp. Fig. 7 A**). This decrease specifically occurred in cardiomyocytes (**Supp. Fig. 7B**). Bulk RNA-seq analysis on cardiac samples obtained from AAV-*Ifng*-treated mice confirmed that the ‘interferon gamma response signature’ was among the top down-regulated gene sets in IFNGR^CM-KO^ mice, compared to Cre^CTRL^, while the ‘inflammatory response’ gene set remained similar between the groups (**Fig. 6 D**). These findings are consistent with specific IFNGR ablation, along with preserved IFN-γ responsiveness in inflammatory cells of the myocardium. Interestingly, the gene sets ‘oxidative phosphorylation’ and ‘fatty acid metabolism’ were among the top up-regulated IFNGR^CM-KO^ mice treated with AAV-*Ifng*, compared to Cre^CTRL^ AAV-*Ifng*-treated controls, suggesting this signature is primarily driven by cardiomyocyte-specific IFN-γ signaling (**Fig. 6 C, E**). These findings indicate that cardiomyocyte-specific IFNGR signaling drives transcriptional reprogramming of cardiac metabolic genes through direct mechanisms.

**Figure 6:**
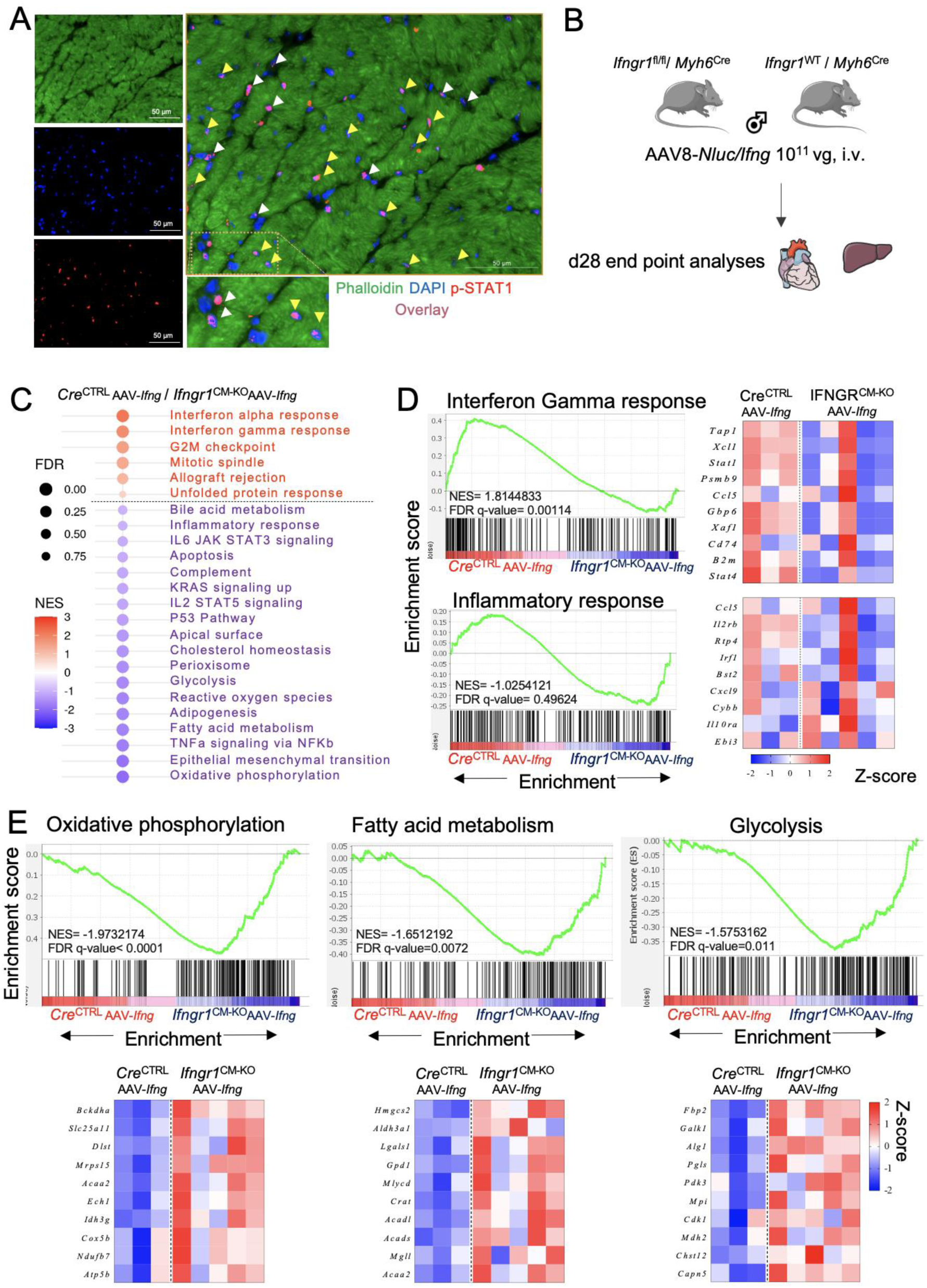
IFN-γ-induced cardiac metabolic rewiring is driven by direct effects in cardiomyocytes. **A**: Immunofluorescence staining of heart sections from AAV-*Ifng*-injected mice: individual channels display phalloidin (green) for myocyte actin filaments, DAPI (blue) for nuclei, and p-STAT1 (red). The overlay on the right highlights p-STAT1 signals, with yellow arrows indicating localization in myocytes and white arrows indicating localization in non-myocytes. **B**: Schematic representation of the cardiomyocyte-specific IFNGR1 knock-out model. **C**: Dot blot showing significantly up- and down-regulated gene sets, as obtained from bulk RNA sequencing data of hearts from the two groups. Color scale represents normalized enrichment score, and circle size shows the false discovery rate (FDR). **D**: Gene set enrichment analysis of interferon gamma response and inflammatory response, with heat map of z-scores for representative top genes, in the heart, among the two groups. **E**: Gene set enrichment analysis of oxidative phosphorylation, fatty acid oxidation, and glycolysis, with heat map of z-scores for representative top genes, in the heart, among the two groups. Blue represents transcript down-regulation while red indicates up-regulation.

## Discussion

IFN-γ is a pro-inflammatory cytokine primarily expressed by NK and T cells commonly up- regulated across several myocardial diseases ^12,14,15,17,24,38^, including autoimmune myocarditis, Chagas cardiomyopathy, pressure-overload induced HF, and MI, as recently reported by independent groups. Previous studies have established that IFN-γ can impact cardiac fibroblasts ^16^, macrophages ^10,39^, and endothelial cells ^40^, ultimately contributing to local inflammation and fibrosis. While its roles in controlling in situ inflammatory mechanisms are well established, the direct impact of IFN-γ on cardiomyocyte function remains incompletely understood. In the present study, we developed an AAV-based model to systemically overexpress IFN-γ and dissect its effects on cardiac metabolic rewiring, independent of other confounding factors. By combining bulk RNA seq, targeted metabolomic analyses, computational modelling, in vivo PET imaging and mitochondrial functional assays in various transgenic mouse models, we found that IFN-γ signaling in cardiomyocytes suppresses oxidative phosphorylation and fatty acid oxidation while enhancing glucose uptake. These findings shed new light on how a cytokine can modulate myocardial function through direct effects on cardiomyocyte metabolism.

A previous study reported that transgenic mice over-expressing IFN-γ developed lethal progressive multiorgan inflammation marked by starkly reduced body weight ^28^. Other independent studies using a transgenic mouse strain constitutively over-expressing IFN-γ, reaching serum concentrations of 20,000-60,000 ρg / ml, observed cardiac fibrosis, ventricular dysfunction, and high mortality rates associated with chronically elevated IFN-γ levels ^7^. In sharp contrast, our model, based on AAV-induced IFN-γ expression rather than on constitutive over-expression, did not cause weight loss, gross histopathological alterations in the heart or liver, or spontaneous mortality. It might therefore provide a suitable system for investigating chronic IFN-γ-driven cardiac inflammation and metabolic rewiring in a more physiological context. Moreover, using an AAV-based expression system offers greater flexibility, as the cytokine expression can be time-controlled, scaled down according to dosing regimens, and adapted for tissue-specific expression based on viral tropism and promoter choice.

While the roles of IFN-γ in promoting cardiac inflammation has been established in previous studies ^16,38^, its relevance in modulating cardiac metabolism remains elusive. At the transcriptional level, we found an inverse correlation between the ‘IFN-γ response signature’ and ‘oxidative phosphorylation’ gene sets. This pattern has been consistently found across multiple available datasets spanning different myocardial conditions ^24^, but the possible causal links between these gene sets remain poorly understood. For instance, in a previous study, we reported a down-regulation of ‘fatty acid oxidation’ and ‘oxidative phosphorylation’ alongside an up-regulation of ‘IFN-γ response’, shifts that are associated with an increased IFN-γ signature from IFN-γ-producing CD8^+^ T cells in aged myocardium ^24,41–50^. Another study on hearts of aged mice revealed transcriptional modifications in several genes involved in mitochondrial function and metabolic enzymes, alterations that primarily suggest defective mitochondrial oxidative potential ^41^. Likewise, similar metabolic profiles are observed during HFrEF in both patients and murine model, which are characterized by a shift from fatty acid oxidation towards glucose ^42–45^.

It has been reported that failing hearts can exhibit changes in metabolic enzyme expression and activity, characterized by decreased fatty acid oxidation and mitochondrial oxidative metabolism, together with elevated glucose uptake and glycolysis ^46^. Moreover, patients with atrial fibrillation exhibit up-regulated IFN-γ response in the atrium, which is associated with down-regulated oxidative phosphorylation ^47^. Consistently, we found increased glycolytic signatures and significantly decreased expression of ATP5B protein from complex V within the electron transport chain ^48^. These results suggest that the mitochondria had functionally reduced oxidative phosphorylation potential shifting energy provision towards the cytosol. These observations recapitulate mitochondrial and energy metabolism alterations observed in patients with Chagas cardiomyopathy, suggesting that cytokine-induced mitochondrial dysfunction might be an important pathophysiological factor in cardiac diseases associated with inflammation ^50^. Similarly, Ashour et al. applied IFN-γ to stimulate iPSC-derived cardiomyocytes and demonstrated that, while neither cellular nor mitochondrial morphology was affected, the mitochondria exhibited defective oxidative phosphorylation ^24^. Another previous study demonstrated that stimulating human ventricular cardiomyocytes with both IFN-γ and TNF resulted in in mitochondrial dysfunction, with reduced fatty acid oxidation and down-regulation of proteins involved in oxidative phosphorylation and the electron transport chain ^49^.

In the present study, we mechanistically addressed causal links between IFN-γ and myocardial mitochondrial respiration pathways using targeted IFN-γ expression and cell-specific ablation studies in vivo. Moreover, we integrate evidence across multiple levels, including transcriptional, metabolic, enzymatic, and functional analyses. Interestingly, these multidimensional metabolic reprogramming events were also reflected in vivo in cardiac fuel uptake, with an IFN-γ-dependent rise in cardiac glucose uptake and up-regulation of glucose transporter expression. These findings conceptually align with previous studies, which have shown in various cardiac disease models that ischemic and failing hearts increasingly rely on glucose metabolism through glycolysis to meet their energy demands ^51,52^.We observed reduced pyruvate dehydrogenase activity, along with lower *Pdha1* and higher *Pdk3* expression, which indicate impaired aerobic glucose oxidation and disrupted pyruvate entry into the Krebs cycle. Further, high levels of aspartate, alanine, and glutamate suggest elevated amino acid catabolism and anaplerosis flux into the Krebs cycle ^53–55^. These metabolic shifts reflect an adaptive metabolic response to sustain energy provision and cardiac function upon IFN-γ-mediated stress in the heart ^56–59^. Interestingly, genetic ablation of IFNGR in cardiomyocytes confirmed that direct IFN-γ signaling in cardiomyocytes drives, at least in part, the observed cardiac metabolic alterations. While it cannot be excluded that IFN-γ-induced cardiac metabolic reprogramming also involves crosstalk with other cardiac cells, including macrophages and fibroblasts that express high levels of IFNGR ^16,20,22,23^, these observations pointing to direct lymphocyte–cardiomyocyte crosstalk provide novel mechanistic insights that may be further explored for therapeutic purposes in the future.

Overall, by integrating immune signaling, metabolic regulation, and cardiac function, our study demonstrates a causal link between IFN-γ signaling in cardiomyocytes and cardiac metabolic rewiring. We identify IFN-γ as a central regulator of cardiomyocyte glucose uptake and glycolysis, demonstrating that, in addition to its classical immune functions, IFN-γ plays a direct role in modulating cardiac metabolism. Given that this cytokine is expressed in a broad range of myocardial conditions, including HF, myocarditis, and MI, our findings provide mechanistic insight into how chronic adaptive immune activation can influence myocardial function, beyond canonical inflammatory pathways.

## Non-standard abbreviations

AAV: Adeno-Associated Virus
ATP5b: ATP Synthase Beta
CD: Cluster of Differentiation
CVDs: Cardiovascular Diseases
FA: Fatty Acid
GAPDH: Glyceraldehyde 3-Phosphate Dehydrogenase
GSEA: Gene Set Enrichment Analysis
HF: Heart Failure
HFpEF: Heart Failure with Preserved Ejection Fraction
IHD: Ischemic Heart Diseases
IFN-γ: Interferon Gamma
IFNGR: Interferon gamma receptor
iPSCs: Induced Pluripotent Stem Cells
LDH: Lactate Dehydrogenase
MHCII: Major Histocompatibility Complex Class II
PD-L1: Programmed Death Ligand 1
PDH: Pyruvate Dehydrogenase
p-STAT1: Phosphorylated Signal Transducer and Activator of Transcription 1
PM: Pyruvate-Malate
TAC: Transverse Aortic Constriction
TNF: Tumor Necrosis Factor.

## Acknowledgements

We thank Lisa Popiolkowski and Elena Vogel for their skillful technical assistance. The figures were prepared with the help of SMART Servier Medical Art.

