## Supplemental Figures for "Interferon gamma signaling drives cardiac metabolic rewiring"

**Supplemental Material**

Supplemental Methods

Supplemental Figures 1-7

### Methods

#### Adeno-associated virus production

HEK 293T (ATCC CRL-1573) cells were maintained on complete DMEM media: DMEM, high glucose (Gibco #11965092), supplemented with 10% FCS, 1x Glutamax (Gibco #35050061), 1x penicillin/streptomycin (Pen/Strep) (Gibco #15070063), 1x Sodium Pyruvate (Gibco #11360070), 1x MEM NEAA (Gibco #11140050), and 25µM 2-mercaptoethanol 50mM (Gibco #31350010). For maintenance, the cells were split 1:8 twice per week. Cells were transfected at 70% confluency and media changed 2-4 hours before transfection to growth DMEM media: DMEM, high glucose (Gibco #11965092), supplemented with 5% FCS, 1x Glutamax (Gibco #35050061), 1x Pen/Strep (Gibco #15070063), and 1x Sodium Pyruvate (Gibco #11360070). Transfection was performed using Polyethyleneimine (PEI) (PolyScience) plasmid triple transfection with pAAV2/8 capsid plasmid (James M. Wilson (Addgene plasmid # 112864 ; http://n2t.net/addgene:112864 ; RRID:Addgene_112864)), pAdDeltaF6 helper plasmid (James M. Wilson (Addgene plasmid # 112867 ; http://n2t.net/addgene:112867 ; RRID:Addgene_112867)), and cargo plasmids designed with Vector builder: pAAV[Exp]-Alb> *mIfng*:T2A:*Nluc*:WPRE (VB220517-1150ajn) or pAAV[Exp]-Alb>*Nluc*:WPRE (VB230130-1072dyr). For viral production, following overnight incubation with growth media containing PEI and plasmids polyplexes, media was changed to production DMEM media: DMEM, low glucose, pyruvate (Gibco # 11885084), supplemented with 1% FCS, 1x Glutamax (Gibco #35050061), 1x Pen/Strep (Gibco #15070063), 10mM HEPES, 1M Buffer Solution (Gibco #15630049), and 0.075% Sodium Bicarbonate 7.5% solution (Gibco #25080060). AAVs were harvested from the culture supernatant on day 3 post-transfection, combined with virus collected from both the cell pellet and supernatant on day 5, and subjected to multiple purification steps. For in vivo-grade quality, purification was performed using iodixanol discontinuous gradient ultracentrifugation (OptiPrep, #1114542). Viral quantification was carried out by qPCR, and quality assessment was performed by western blotting, following established AAV production protocol [1].

#### In vitro validation

Transduction efficiency was checked on AML12 hepatocyte cell line, ATCC CRL-2254 [2]. Cells were maintained on complete DMEM/F-12 media: DMEM/F-12, no phenol red (Gibco #21041025), 10% FCS, 1x Pen/Strep (Gibco #15070063), 1x Insulin-Transferrin-Selenium (ITS -G) (Gibco #41400045), and 4 ng/ml dexamethasone (Sigma-Aldrich D4902-25mg). For maintenance, the cells were split 1:4 twice per week. Cells were transduced with AAVs at 70% confluency using $10^{10}$, $10^{9}$, and $10^{8}$ viral genome load per $10^{6}$ cells, and incubated overnight.

Medium was changed the next day to low protein DMEM/F-12 (1% FCS), and gently replaced again three days post-transduction. Six days post transduction, luciferase activity (Nano-Glo luciferase assay system N1130) and IFN-γ ELISA measurement were preformed from cell culture supernatant (Mouse IFNg ELISA MAX standard set ,430801) according to manufacturer's instructions. Plates were recorded using Tecan Spark Multimode Microplate Reader.

**Animals**

C57BL/6J male mice six to eight weeks old were imported from the Jackson Laboratory and housed under specific pathogen-free (SPF) conditions, with a controlled light–dark cycle and a standard diet. B6.FVB-Tg(*Myh6*-cre)2182Mds/J mice were crossed with C57BL/6N-*Ifngr1*$^{tm1.1Rds}$/J mice (both imported from Jackson Laboratory. Strain#: 011038 and Strain#: 025394 respectively) to generate C57BL/6N-*Ifngr1*$^{tm1.1Rds}$/J Tg(*Myh6*-cre)2182Mds/J (*Myh6*$^{Cre}$ *Ifngr1*$^{fl/fl}$) transgenic mice with cardiomyocyte-specific IFNGR1 knockout. No experimental animals enrolled in the study were excluded from analysis. All animal procedures were approved by the local authorities (Regierung von Unterfranken) and conformed to the guidelines from Directive 2010/63/EU of the European Parliament on the protection of animals used for scientific purposes [3].

**Systemic IFN-γ over expression in vivo**

C57BL/6J mice were intravenously injected via the tail vein by $10^{11}$ viral genomes (vg) per mouse of AAV, cardiac function was monitored by echocardiography at day 14 and day 28 post injection. All animals were sacrificed 28 days after viral injection by cervical dislocation and then perfused with PBS-heparin (50 IU/ml) to flush the coronary circulation and remove blood-borne leukocytes, according to a described protocol [4] and organs were collected for end point analyses. The choice of day 28 as the endpoint is based on AAV viral kinetics, as AAVs typically become detectable one week after tail vein injection, with peak expression occurring between two and six weeks [5,6].

**Echocardiography**

Echocardiography measurements were performed with a Vevo 1100 instrument (VisualSonics, Amsterdam, Netherlands) with a MS 400 echo transducer coupled to a 30-MHz probe developed for mice studies. Briefly, mice were maintained under slight isoflurane anesthesia (0.5-1.5% vol/vol $O_2$) on a heated bed (39°C), and images were acquired on the short axis at apex, midpapillary (B and M mode) and basal (only B mode) levels and on the long axis (B and M mode) as well as the 4-chamber view. Basal heart rate between 450-600 bpm was targeted during assessment. All assessments were performed by an experienced scientist. The analyses were performed blinded using the manufacturer's software (Vevo® LAB 1.7.1).

### [$^{18}$F]FDG PET imaging

As recommended by Ribeiro et al [7], animals were fasted in clean cages without access to food and were provided with water ad libitum for 12 hours prior to [$^{18}$F]FDG PET. During the fasting period temperature was maintained constant in an IVC System (Smart Flow Tecniplast Germany) and additional nesting material and enrichment was provided. To prevent the effect of isoflurane in myocardial glucose metabolism, we used conscious [$^{18}$F]FDG administration and 60 minutes uptake followed by unconscious PET scanning as recommended for [$^{18}$F]FDG cardiac imaging. In concrete, after the fasting period , the animals were removed briefly from the IVC cages to receive [$^{18}$F]FDG intraperitoneal injection (5 MBq). Then the animals were placed back to the IVC cage system with controlled temperature. Fifty five minutes after [$^{18}$F]FDG injection, the animals were set under isoflurane anesthesia and placed in the heated bed of a dedicated micro PET scanner (Inveon, Siemens Medical Solutions). Image acquisition time was 10 minutes. After data acquisition, animals return to their original home cages with fresh food. Cardiac [$^{18}$F]FDG uptake in vivo was quantified as the percentage of the initially injected dose per cubic centimeter and was used to calculate the heart to mediastinum ratio (HMR) [7]. To confirm adequate fasting, animals were weighed before and after fasting. There was no difference in weight loss related to the treatment. Blood glucose was controlled after isoflurane anesthesia and was not different among the genotypes.

### [$^{18}$F]FTHA Biodistribution

In preparation for [$^{18}$F]FTHA biodistribution, the animals were set for fasting in clean cages without food and water was provided ad libitum for 12 hours. During the fasting period temperature was maintained constant and additional nesting material and enrichment was provided. Animals were set under isoflurane anesthesia (2%) in a heated bed and the tracer (0.5MBq) was administer via a tail vein catheter. Sixty minutes post [$^{18}$F]FTHA administration the organs of interest were harvested and weighed. Radioactivity of each organ was measured in a γ-counter (Gamma Counter Wizard2 2480 Perkin- Elmer LAS). Results are expressed as the percentage of injected dose per gram of tissue (%ID/g), adjusted for decay and normalized to organ weight. Finally, organ-to-blood ratios were calculated by dividing %ID/g of organ to the corresponding blood of the animals. To confirm adequate fasting, animals were weighed before and after fasting. There was no difference in weight loss related to the treatment. Serum triglycerides level was controlled during organ collection and was not different among the genotypes.

**Cardiomyocytes isolation**

After sacrificing the mice, hearts were rapidly cannulated and retrogradely perfused with a liberase/trypsin solution (Roche Diagnostics GmbH; protocol PP00000125 from The Alliance for Cellular Signaling). The resulting single cell suspension was centrifuged at 100 × g for 60 s at 4 °C to pellet rod-shaped cardiac myocytes. The resulting supernatant was transferred to a new 15-ml tube and centrifuged at 500 ×g for 5 min at 4 °C to obtain the non-myocyte fraction [8]. Both cardiomyocyte and non-cardiomyocyte pellets were snap-frozen in liquid nitrogen.

**Protein and RNA extraction**

Heart, liver, Lung and skeletal muscles were homogenized in PBS (#L 182-50) using Qiagen tissue homogenizer and disposable probes (Qiagen #990890) for luciferase activity measurements, or in RLT supplemented with β-mercaptoethanol lysis buffer for protein and RNA extraction using Qiagen all prep kit (#80004) according to manufacturers' instructions. RNA concentration and purity were routinely checked using NanoDrop 2000 (Thermo Scientific™ ND2000). Protein preparations were resuspended in 5% SDS in distilled water, and total protein concentration was checked by Pierce™ BCA Protein Assay Kit (#23227).

**Bulk RNA sequencing**

RNA quality was assessed using a 2100 Bioanalyzer with the RNA 6000 Nano kit (Agilent Technologies), ensuring all samples had an RNA integrity number (RIN) ≥7.5. DNA libraries suitable for sequencing were prepared from 200 ng of total RNA using oligo-dT capture beads for poly-A enrichment, following the TruSeq Stranded mRNA Library Preparation Kit (Illumina) protocol at half volume. After 15 cycles of PCR amplification, the size distribution of the barcoded DNA libraries was estimated to be ~310-320 bp via electrophoresis on Agilent DNA 1000 Bioanalyzer microfluidic chips. Sequencing of pooled libraries, spiked with 1% PhiX control library, was conducted at 25 million reads per sample in single-end mode with a 100 nt read length on the NextSeq 2000 platform (Illumina). Demultiplexed FASTQ files were generated using bcl-convert Version 4.0.3 and 4.3.6 (Illumina). To ensure high sequence quality, Illumina reads were quality- and adapter-trimmed using Cutadapt (Martin, 2011) version 2.5 with a cutoff Phred score of 20 in NextSeq mode, discarding reads without any remaining bases (command line parameters: --nextseq-trim=20 -m 1 -a AGATCGGAAGAGCACACGTCTGAACTCCAGTCAC). Processed reads were subsequently mapped to the mouse genome (GRCm39) using STAR v2.7.2b with default parameters [9]. Read counts at the exon level were summarized for each gene using featureCounts v1.6.4 from the Subread package. Multi-mapping and multi-overlapping reads were counted strand-specifically and reversely stranded with fractional counts for each alignment and overlapping feature (command line parameters: -s 2 -t exon -M -O --fraction). The count output was used to identify differentially expressed genes using DESeq2 [10] version 1.24.0. Read counts were normalized by DESeq2, and fold-change shrinkage was applied by setting the parameter “betaPrior=TRUE”. Differential gene expression was determined at an adjusted p-value (padj) after Benjamini-Hochberg correction < 0.05 and |log2FoldChange| ≥ 1.

**Reverse Transcription Quantitative Polymerase Chain Reaction (RT-qPCR)**

Adeno-associated virus titre quantitation was performed as previously described [1]. Briefly, extra-viral DNA was digested using DNase I (Sigma-Aldrich AMPD1-1KT) by incubating the viral suspension at 37 °C for 15 minutes. Viral DNA was then released by adding 2× capsid lysis buffer and heating at 95 °C for 10 minutes, followed by neutralization with Tris-HCl (pH 8.0) and a 50-fold dilution. qPCR was performed using Fast SYBR™ Green Master Mix (Applied biosystems #4385612) with ITR-specific primers; 100 nM of ITR-Fwd (forward primer for ITR) (5’-GGAACCCCTAGTGATGGAGTT-3’), 340 nM of ITR-Rev (reverse primer for ITR) (5’-CGGCCTCAGTGAGCGA-3’) and a standard curve generated from 5-fold serial dilutions of a plasmid containing ITR sequences. PCR was performed under the following conditions: 50 ºC for 2 minutes, 95 ºC for 2 minutes, followed by 40 cycles at 95 ºC for 3 seconds, and annealing/extension at 60 ºC for 30 seconds. The final titer of the sample, expressed in viral genomes per microliter (vg/µl), was obtained by multiplying the quantified viral genome copies by 2500, based on interpolation in the standard curve.

*Ifng* (Mm01168134_m1) and *Ifngr1* (Mm00599890_m1) mRNA expression levels were quantified in different tissues collected from in vivo experiments by qPCR using TaqMan universal PCR master mix (#4304437). Relative expression was normalized to 18S rRNA (Hs99999901_s1) as an endogenous control. Reactions were performed on Bio-Rad CFX96 Touch real time-PCR-System.

**Western blotting**

AAV quality and purity were checked by staining the blots for total protein of the viral suspension using colloidal gold stain (Bio-rad #1706527) and checked for three distinct AAV bands VP1 90 kDa, VP2 72 kDa, and VP3 63 kDa.

p-STAT1 were checked using Phospho-Stat1 rabbit mAb (Cell signalling #9167), IFNGR1 rabbit mAb (Cell signalling #84318) and respiratory chain complex V using ATP5B primary antibody (Antibodies.com A14964) then the membrane was treated with Goat anti-Rabbit IgG (H+L) Cross-Adsorbed Secondary Antibody, HRP (Invitrogen #G-21234), while GAPDH Mouse mAb (Cell signalling #97166) primary antibody was subsequently treated with anti-mouse IgG, HRP-linked Antibody (Cell signalling #7076). Finally, specific binding was detected using Pierce ECL Western Blotting Substrate (#32106) applied to the membrane. Visualization of developed protein bands was done using the ChemiDocTM Imager (Bio-Rad) which were further quantified using the software Image Lab 6.0 (Bio-Rad).

For sequential detection of multiple proteins, membranes were stripped twice for 10 min with stripping buffer (1.5% glycine, 0.1% SDS, 1% Tween-20, pH 2.2) under shaking, washed with distilled water followed by TBST, re-blocked, and subsequently reprobed with the different antibodies.

**Gas Chromatography-Mass Spectrometry (GC-MS) Polar Metabolite Analysis**

Metabolites were isolated as described previously [11]. Briefly, frozen heart tissue samples were homogenized in 0.4 mL −80 °C cold MS-grade methanol, acetonitrile, and water at a volume ratio of 40:40:20 (v/v/v/). Samples were freeze-thawed three times to achieve further disruption of the tissue sample. Extracts were centrifuged at 13,000 rpm for 5 min at 4 °C. The supernatant was transferred to clean glass tubes, and D27-myristic acid (0.15 µg/µL, Cat. No. 366889, Sigma-Aldrich; St Louis, MO) was added as an internal run standard. Vials were covered with a breathable membrane, and metabolites were evaporated to dryness under vacuum. Dried samples were stored at -80°C for further GC-MS analysis. Dried metabolites were dissolved in 20 µL of Methoxamine (MOX) Reagent (2% solution of methoxyamine-hydrogen chloride in pyridine; Cat. No. 89803 and 270970; Sigma-Aldrich; St Louis, MO) and incubated at 30°C for 90 minutes. Subsequently, 90 µL of trimethylsilyl (TMS) was added, and samples were incubated under shaking at 37°C for 30 minutes. GC-MS analysis was conducted using an Agilent 8890 GC coupled with an Agilent 5977-mass selective detector. Metabolites were separated on an Agilent HP-5MS Ultra Inert capillary column (Cat. No.

19091S-433UI). For each sample, 1 µL was injected at 250°C using helium gas as a carrier with a flow rate of 1.1064 mL/min. For the measurement of polar metabolites, the GC oven temperature was kept at 60°C and increased to 325°C at a rate of 10°C/min (10 min hold), followed by a post-run temperature at 325°C for 1 min. The total runtime was 37 minutes. The MS source and quadrupole were kept at 230°C and 150°C, respectively, and the detector was run in scanning mode, recording ion abundance within 50 to 650 m/z. Metabolomic spectra were annotated in the Agilent Masshunter software by matching features to NIST and Fiehn RTL library. Metabolite annotations were made based on mass errors for all qualifier annotations to library values, GC retention times were within 6 s of elution times from library features. The list of metabolites and corresponding peak abundances was then statistically evaluated.

#### Flux balance analysis using CardioNet

*In silico* simulations were conducted using the metabolic network of the mammalian heart metabolism, CardioNet.[11-13] Metabolite abundances from GC-MS analysis and enzymatic activities were used as constraints for network reactions and integrated for flux balance analysis (FBA) to determine flux distributions ($v_m$) as described previously [12,14-16]. Objective functions were defined to maximize ATP hydrolysis and biomass synthesis [11,14,15,17]. FBA was applied to identify steady-state flux distributions that agree with applied substrate uptake and release rates and changes in quantified metabolite pools. The GUROBI LP solver was used to calculate optimal solutions for each FBA problem [18].

#### Enzyme activity colorimetric assays

The activity of phosphofructokinase (PFK), lactate dehydrogenase (LDH) and pyruvate dehydrogenase (PDH) were monitored using commercially available colorimetric kits (MAK093, MAK006 and MAK183, Sigma-Aldrich), following the manufacturer's instructions.

#### Histological staining

Heart and liver samples were cryopreserved in Tissue Tek® Optimal Cutting Temperature compound™ and the histological slices (10µm) were sliced using cryostat Leica CM1850 and stained by Masson's trichrome staining (Polyscience # 25088) according to manufactures instructions. Briefly, all slides were fixed in 10% neutral buffered formalin at room temperature for 1 hour then incubated in Bouin's solution overnight at room temperature. Then slides were gently washed in running tap water for 5 minutes to remove the picric acid. Slides were next stained in Weigert's Iron Hematoxylin working solution for 10 minutes. Then slides were rinsed in distilled water followed by staining in Biebrich Scarlet-Acid Fuchsin solution for 5 minutes, rinsed, and incubated in Phosphotungstic/Phosphomolybdic acid for 10 minutes. Slides were then stained in Aniline Blue for 5 minutes, rinsed three times in distilled water, and immersed in 1% acetic acid for 1 minute. The slides were dehydrated using 50% ethanol, 75% ethanol, followed by 100% ethanol, for 2 minutes each then cleared in Xylol:ethanol 1:1, followed by Xylol for 5 minutes each and finally mounted using entellan (Sigma-Aldrich #1079600500).

Immunofluorescence staining of heart sections (7µm) was performed for p-STAT1 (cell signalling #9167) and secondary anti rabbit Alexa Fluor plus 555 (Invitrogen #15636746), Phalloidin Alexa Fluor 647 (Invitrogen # A22287), and DAPI (Invitrogen # D1306). Briefly, the slides were permeabilized with ice-cold 100% methanol for 10 minutes at –20°C then rinsed in PBS for 5 minutes. The specimen was blocked in Blocking Buffer (1X PBS / 1X Carbo-free blocking solution / 0.3% Triton™ X-100) for 60 minutes. The primary antibody (in antibody dilution Buffer (1X PBS / 1% BSA / 0.3% Triton X-100) was then applied and incubated overnight at 4°C. The slides were rinsed in PBS three times 5 min each, and then incubated in fluorochrome-conjugated secondary antibody diluted in antibody dilution buffer for 2 hours at room temperature in the dark. Thirty minutes before the end of incubation of the secondary antibody, Phalloidin was added according to dilution instructions of manufacturers, generally, with final concentration of 1 µg/ml. Finally, DAPI was added to all the spots and the slides were further incubated for 10 min. Next, all the slides were mounted with Mowiol.

All slides were imaged using Compact Fluorescence Microscope BZ-X800.

**Flow cytometry analysis**

Mice were injected intravenously with an anti-CD45.2 antibody (4 µg per mouse, clone 104) 3 minutes prior to euthanasia, to further eliminate potentially contaminating circulating leukocytes. After harvesting the organs, hearts were digested using a modified protocol from Skelly et al 34, using type II collagenase (1000 IU/ml, Worthington), Lungs using Liberase (Roche #05401020001, 0.2 mg/ml) and DNase and finally liver using collagenase type D (Roche # 11088858001, 1mg/ml) and DNase. Digested organs and lymph nodes were gently filtered through a 30 µm mesh (Miltenyi Biotec, Bergisch Gladbach, Germany). Surface staining was performed in the presence of FC-blocking antibody (anti-CD16/CD32, clone 2.4G2, BD Pharmingen) using the following conjugated antibody clones commercially acquired from BioLegend (San Diego, CA, USA) unless stated otherwise: anti-TCRβ (clone H57-597), anti-CD45 (clone 30-F11), anti-CD45.2 (clone 104), anti-CD29 (clone HMb1-1, eBioscience/Thermo Fisher Scientific, Waltham, MA, USA), anti-IA/IE (clone M5/114.15.2), anti-CD31 (clone 390), anti-PD-1 (clone J43, eBioscience/Thermo Fisher Scientific), anti-CD11c (clone N418), anti-CCR2 (clone 475301, R&D Systems, Minneapolis, MN, USA), anti-PD-L1 (clone 10F.9G2), anti-CXCR3 (clone CXCR3-173), anti-CD4 (clone RM4-5), anti-CD8a (clone 65-6.7), anti-CD69 (clone H1.2F3), anti-CD64 (clone X54-5/7.1), anti-Ly6C (clone HK1.4), anti-Ly6G (clone 1A8), anti-CD11b (clone M1/70), and anti-MEFSK4 (clone MEFSK4, Miltenyi Biotec, Bergisch Gladbach, Germany). The optimal antibody concentrations were titrated in our laboratory. In addition, Zombie Aqua™ Fixable Viability Kit (BioLegend) was used for live/dead discrimination (1:1000). Flow cytometry measurements were performed on Attune-NxT (Thermo Scientific, Darmstadt, Germany). Flow cytometry data were analyzed using FlowJo (FlowJo LLC, Ashland, OR, USA). Compensation for spectral overlap was conducted based on single staining controls and, whenever meaningful, the flow cytometry gates were set based on 'fluorescence minus one' (FMO) controls.

**Isolated mitochondrial measurements**

Freshly isolated cardiac mitochondria were used for high-resolution respirometry measurements that were performed as described previously [19]. Oxygen consumption was assayed at 37°C with an Oxygraph-2k high-resolution respirometer and DatLab software was used for data acquisition and analysis (Oroboros Instruments, Innsbruck, Austria). Two mitochondrial respiration protocols were used to quantify oxygen consumption rate upon supplementation of pyruvate (for carbohydrate metabolism) or fatty acids (for β-oxidation) as a fuel. In the "carbohydrate" protocol, measurements of complex I and II activity were performed with 5 mM Na-pyruvate and 5 mM Na-malate or 10 mM succinate, respectively. The metabolites were added as reduced substrates after initially recording residual oxygen consumption resulting in leak respiration, followed by increasing concentrations of ADP (0.03, 0.1, 0.3, 1 mM). For succinate-dependent respiration, complex I was inhibited by 0.5 µM rotenone to prevent reverse electron flux. Finally, oxygen consumption coupled to ADP phosphorylation was inhibited by adding 1.25 µM oligomycin followed by titration with 10 µM DNP to determine uncoupled respiration state 3u. In the "fatty acid" protocol, respiration was measured using 1 mM carnitine, 3 mM malate,10 µM palmitoyl-CoA, and 10 µM oleoyl-L-carnitine. Similar to the "carbohydrate" protocol, increasing amounts of ADP, 1.25 µM oligomycin or 10 µM DNP were added. Mitochondrial membrane potential was simultaneously probed using 1 µM TMRM and Smart Fluo-Sensor Green as described before [19]. $H_2O_2$ flux was measured simultaneously with respirometry in the O2k-Fluorometer using the Amplex Ultra Red-horseradish peroxidase (HRP) system. Also, for these, two protocols were used: one with 5 mM Na-pyruvate and 5 mM Na-malate and another one with the fatty acid mixture as substrates as described above. After addition of substrates, 1 mM ADP and 1.5 µM antimycin A as complex III inhibitor were added. A calibration curve was generated daily using successive additions of known [$H_2O_2$] in the absence of isolated mitochondria.

**Plasma measurements**

Blood was collected into heparin-washed syringes and centrifuged at 4°C for 10 minutes to separate plasma. Plasma levels of IFN-γ were measured using the Mouse IFN-γ ELISA MAX Standard Set (#430801), insulin using the Ultra-Sensitive Mouse Insulin ELISA Kit (#90080), and triglycerides using the EnzyChrom™ Triglyceride Assay Kit (#ETGA-200), according to the manufacturer's instructions.

**Statistical analyses**

The results are shown as the mean ± the standard deviation of the mean (SD) along with the distribution of all individual values in each group. Sample sizes for each group are described in figure legends. Graphs and statistical analyses were performed with GraphPad Prism (version 7.0, GraphPad Software, San Diego, CA, USA). Unpaired two-tailed t-test was used to compare two groups with data following normal distribution. For multiple comparisons between more than two groups, one or two-way analyses of variance (ANOVA) were conducted followed by post hoc test. Differences were considered significant for p values below 0.05.

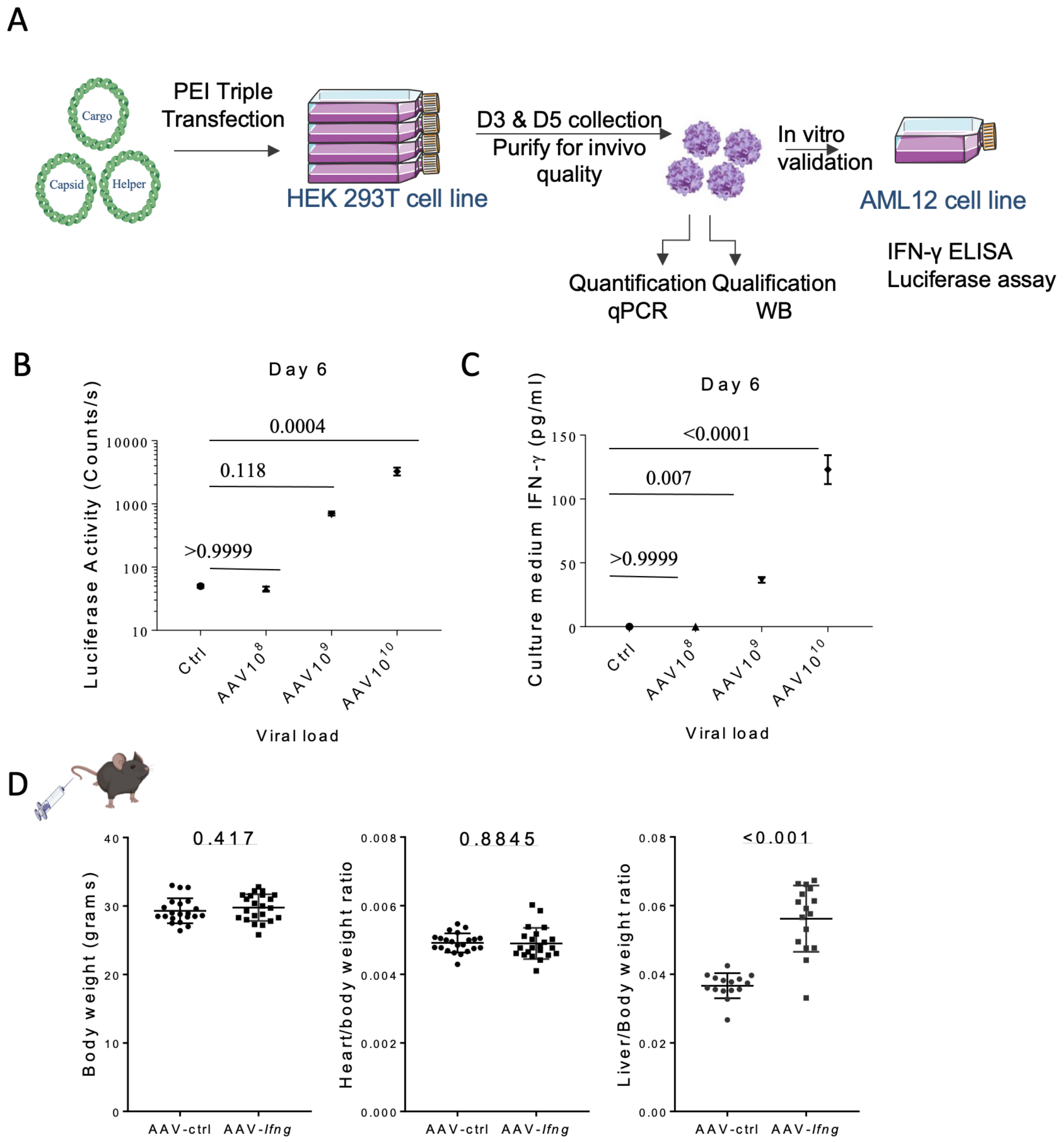


**Supp. Fig. 1: Adeno-associated virus production and in vitro validation. A:** Schematic representation of AAV production followed by in-vitro validation. **B:** Day 6 luciferase activity of AML12 cells transduced with different viral loads against control non- transduced cells. **C:** Day 6 IFN-γ concentration in cell culture medium of AML12 cells transduced with different viral loads against control non- transduced cells. **D:** Graphs comparing post-AAV injection total body weight, heart weight to body weight, liver weight to body weight (from left to right) between AAV-*Ifng* and AAV-ctrl groups. Statistical analysis for graph B, C using one-way ANOVA with multiple comparisons comparing mean of each column to the Ctrl column using Dunnett test, and statistical analysis for graph E using t-test. Graphs B, C, D are scatter plot showing Mean ± SD. p value is written for each comparison.

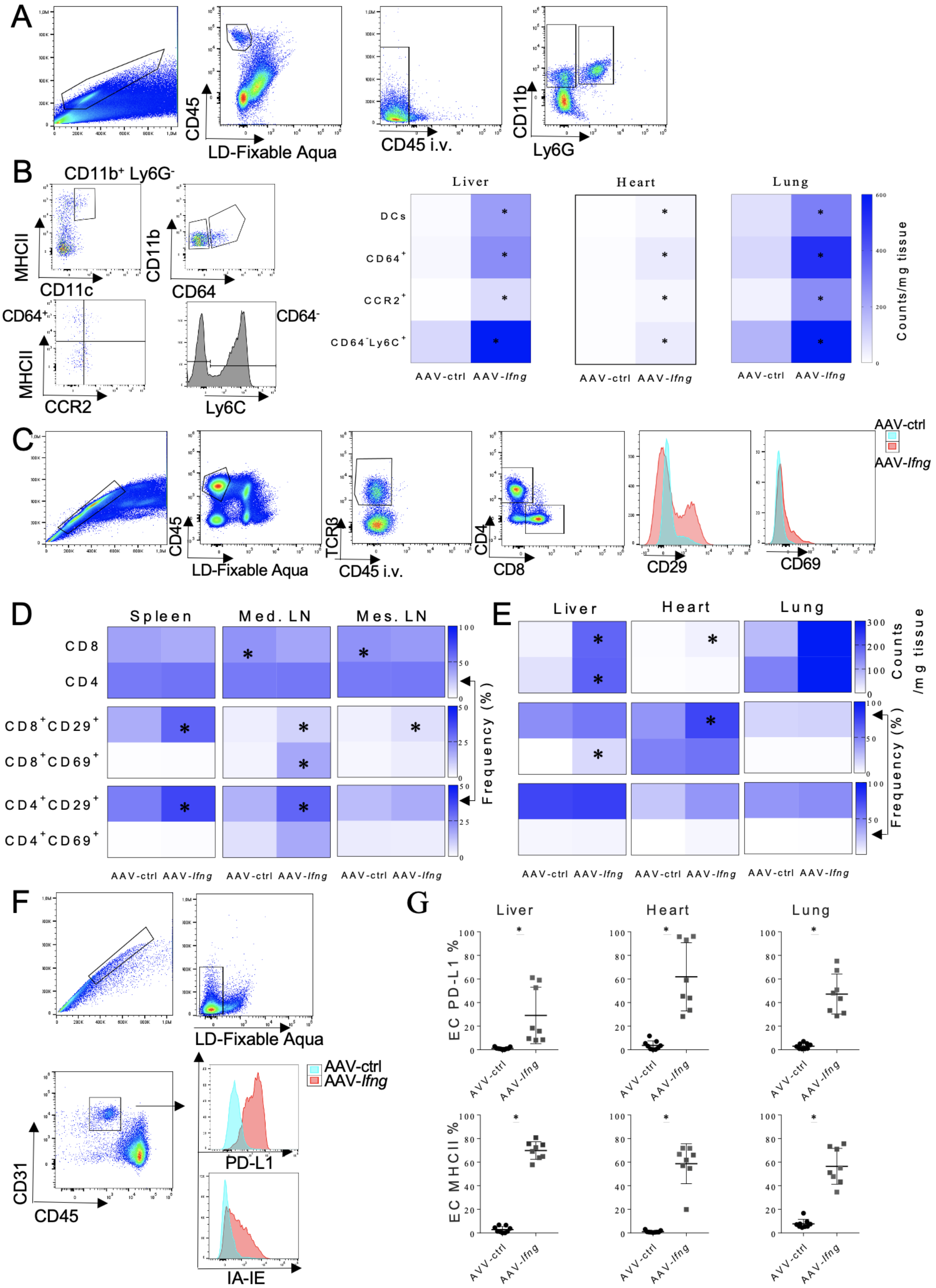


**Supp. Fig. 2: Flow cytometry gating strategy. A:** Flow cytometry gating of myeloid cells. **B:** Heat map quantification of Dendritic cells, Macrophages, Monocyte derived macrophages, and Ly6C $^{hi}$ monocytic cells. Statistical analysis: Multiple t-test. * indicates significance with p value < 0.05. Graphs B are scatter plot showing Mean ± SD. **C:** Flow cytometry gating of T cells and histograms of CD29 and CD69 expression on T cells between the two groups. **D:** Heat map quantification of T cells in spleen, mediastinal lymph node, and mesenteric lymph node. **E:** Heat map quantification of T cells in liver, heart and lung of the two groups. **F:** Flow cytometry gating of cardiac endothelial cells ($CD45^-$ $CD31^+$). **G:** Quantification of MHCII and PD-L1 expression on endothelial cells across various tissues. Statistical analysis using t-test. * Indicates significance with p value < 0.05.

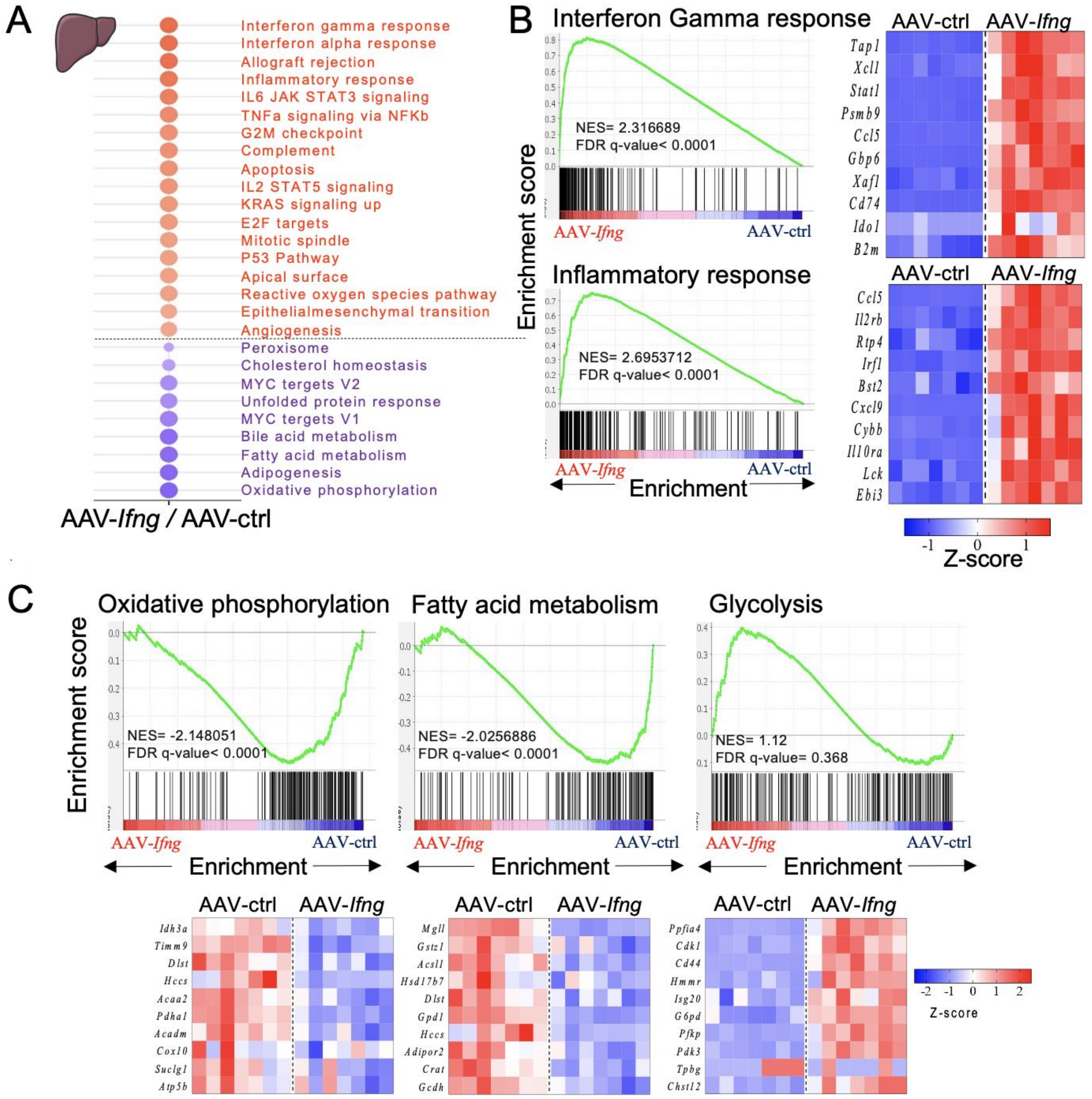


**Supp. Fig. 3: IFN-γ impacts liver transcriptome profile. A:** Dot blot showing significant up and down-regulated gene sets obtained from bulk RNA sequencing data of liver of the two group. Colour scale represents the normalized enrichment score and circle size represents the false discovery rate (FDR). **B:** Gene set enrichment analysis of interferon gamma response and inflammatory response with heat map of z-scores of representative top genes between the two groups in liver. **C:** Gene set enrichment analysis of oxidative phosphorylation, fatty acid oxidation and glycolysis with heat map of z-scores of representative top genes between the two groups in liver. Blue colour represents downregulation while red colour indicates upregulation of transcripts.

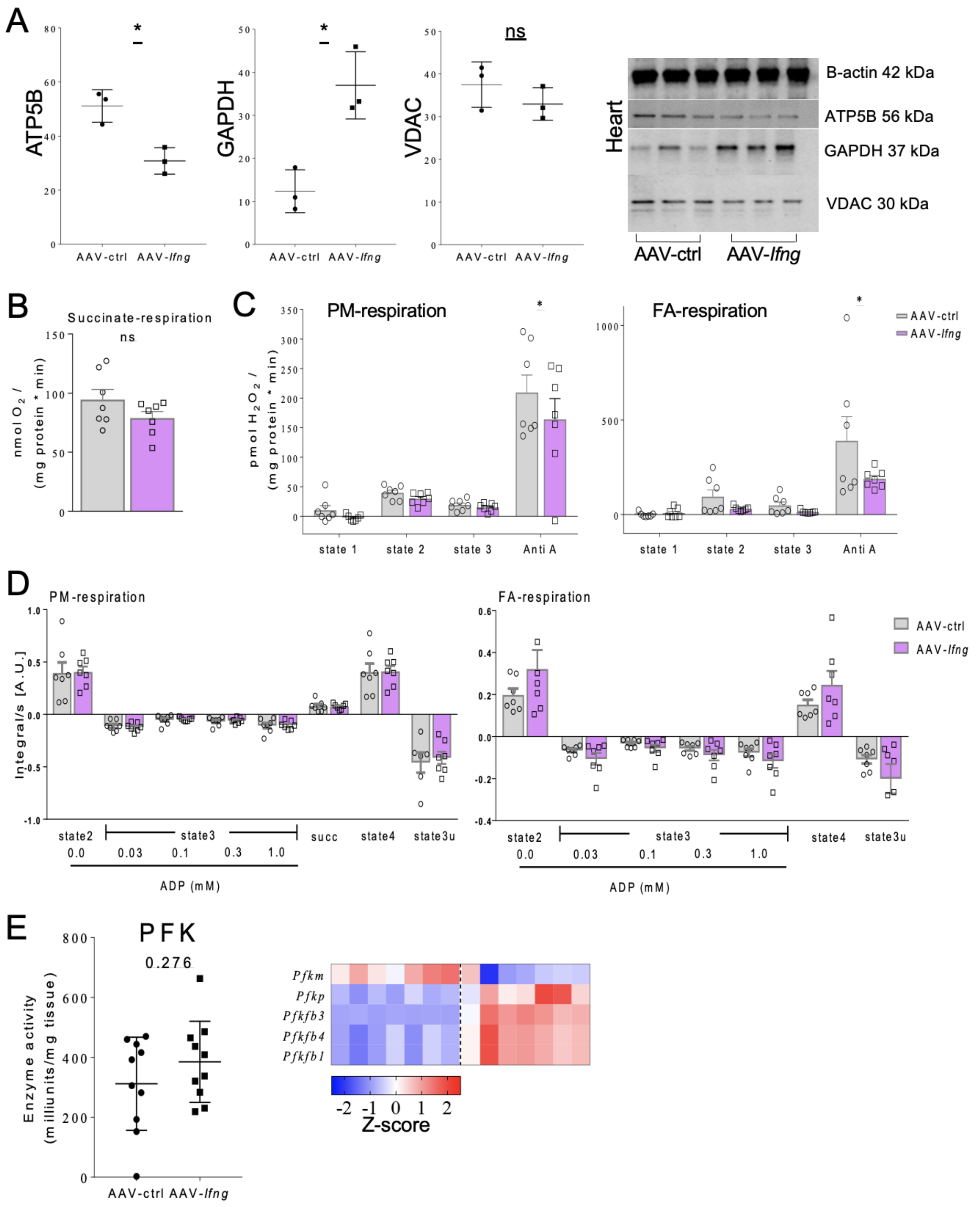


**Supp. Fig. 4: Isolated mitochondria function measurements. A:** Western blotting membrane showing β-actin, ATP5B, GAPDH and VDAC in hearts of the two groups (3 different mice per each group) with quantification graphs of ATP5B, GAPDH and VDAC. **B:** Succinate respiration between the two groups. **C:** Quantification of $H_2O_2$ production on both Pyruvate-Malate dependant respiration and fatty acid complex dependant respiration on different respiration states and with actimycinA (antiA) blocking of complex 3. **D:** Mitochondrial membrane potential measurements using TMRM dye with both PM standard protocol on the left and FA standard protocol on the right for different experimental respiratory states. E: Phosphofructokinase enzyme activity quantification graph between the two groups and heat map of z-score of enzyme genes. Statistical analysis using t-test. * Indicates significance with $p$ value $< 0.05$. Graphs A, B, C, D, E, F are scatter plot showing Mean ± SD.

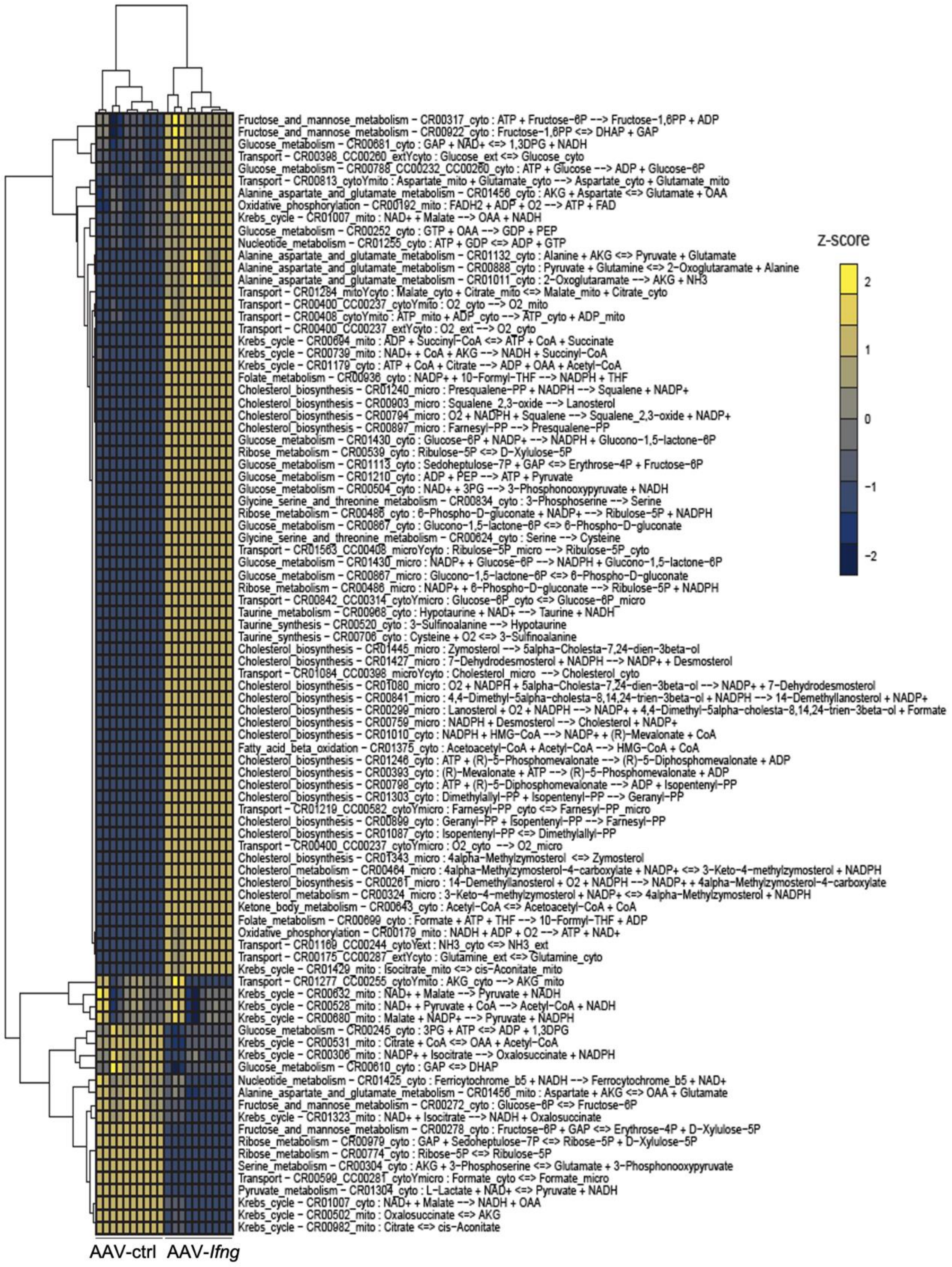


**Supp. Fig. 5:** Heatmap representing unsupervised hierarchical cluster analysis of flux balance analysis with an adjusted p-value <0.01. Experimental data from AAV-ctrl- and AAV-*Ifng*-injected mice was integrated into CardioNet simulations, and metabolic flux rates were calculated using an objective function demanding energy provision and biomass synthesis. Calculated flux rates were compared using a linear regression model and annotated to CardioNet pathways.

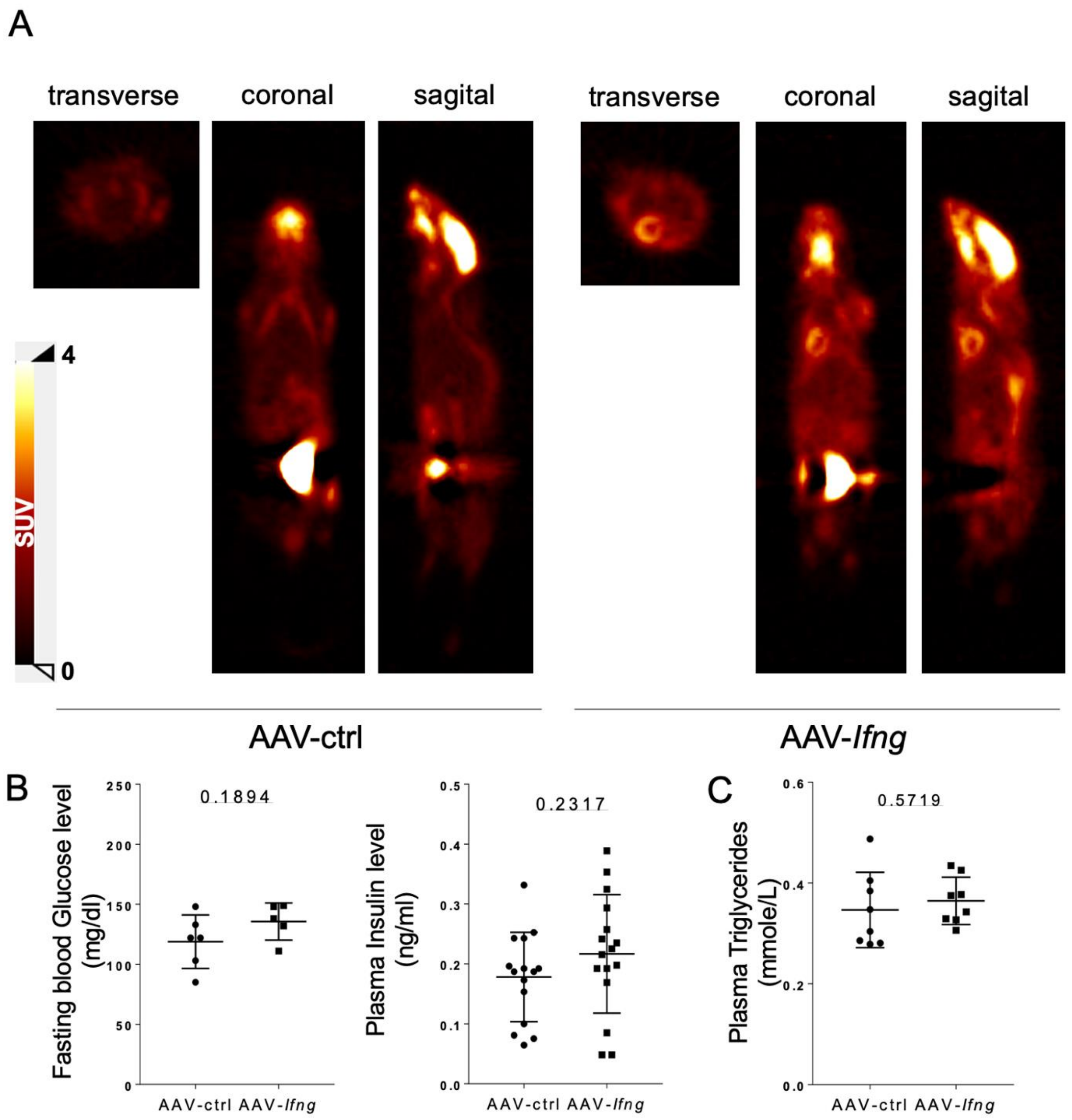


**Supp. Fig. 6: [$^{18}$F]FDG glucose uptake PET imaging. A:** Representative [$^{18}$F]FDG uptake PET images from transverse, coronal, and sagittal planes of the same animal in both AAV-ctrl and AAV-*Ifng*-injected mice. **B:** Fasting blood glucose level, and serum insulin level. **C:** plasma triglycerides level. Data are shown as scatter plot showing mean ± SD and statistical significance were measures by unpaired t test. p value is written for each comparison.

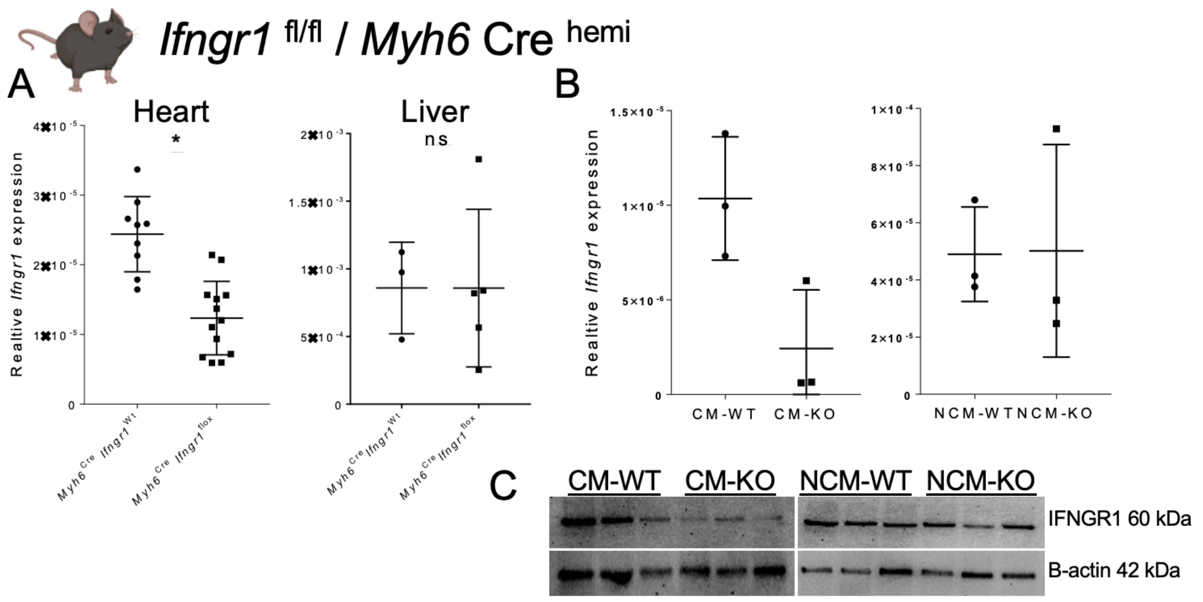


**Supp. Fig. 7: *Ifngr1* $^{fl/fl}$ / *Myh6* Cre $^{hemi}$ transgenic mouse model validation.** A: Quantification graphs of relative *Ifngr1* mRNA expression between *Myh6*$^{Cre}$ *Ifngr*1$^{flox}$ knock out mice group and *Myh6*$^{Cre}$ *Ifngr*1$^{Wt}$ control mice group in heart and liver. B: (on the left) Quantification graph of relative *Ifngr1* mRNA expression between isolated cardiomyocytes of wild type mice and cardiomyocytes of *Myh6*$^{Cre}$ *Ifngr*1$^{flox}$ knock out mice, (on the right) quantification graph of relative *Ifngr1* mRNA expression between non-cardiomyocytes of wild type mice and non-cardiomyocytes of *Myh6*$^{Cre}$ *Ifngr*1$^{flox}$ knock out mice. C: Western blotting membrane showing β-actin and IFNGR1 expression in cardiomyocytes and non-cardiomyocytes of wild type and *Myh6*$^{Cre}$ *Ifngr*1$^{flox}$ knock out mice (3 mice per each group). Statistical analysis using t-test. * Indicates significance with $p$ value $< 0.05$. Graphs A, B are scatter plot showing Mean ± SD
